# Polarized Desmosome and Hemidesmosome Shedding via Exosomes is an Early Indicator of Outer Blood-Retina Barrier Dysfunction

**DOI:** 10.1101/2023.06.12.544677

**Authors:** Belinda J. Hernandez, Nikolai P. Skiba, Karolina Plößl, Madison Strain, Daniel Grigsby, Una Kelly, Martha A. Cady, Vikram Manocha, Arvydas Maminishkis, TeddiJo Watkins, Sheldon S. Miller, Allison Ashley-Koch, W. Daniel Stamer, Bernhard H.F. Weber, Catherine Bowes Rickman, Mikael Klingeborn

## Abstract

The retinal pigmented epithelium (RPE) constitutes the outer blood-retinal barrier, enables photoreceptor function of the eye, and is constantly exposed to oxidative stress. As such, dysfunction of the RPE underlies pathology leading to development of age-related macular degeneration (AMD), the leading cause of vision loss among the elderly in industrialized nations. A major responsibility of the RPE is to process photoreceptor outer segments, which relies on the proper functioning of its endocytic pathways and endosomal trafficking. Exosomes and other extracellular vesicles from RPE are an essential part of these pathways and may be early indicators of cellular stress. To test the role of exosomes that may underlie the early stages of AMD, we used a polarized primary RPE cell culture model under chronic subtoxic oxidative stress. Unbiased proteomic analyses of highly purified basolateral exosomes from oxidatively stressed RPE cultures revealed changes in proteins involved in epithelial barrier integrity. There were also significant changes in proteins accumulating in the basal-side sub-RPE extracellular matrix during oxidative stress, that could be prevented with an inhibitor of exosome release. Thus, chronic subtoxic oxidative stress in primary RPE cultures induces changes in exosome content, including basal-side specific desmosome and hemidesmosome shedding via exosomes. These findings provide novel biomarkers of early cellular dysfunction and opportunity for therapeutic intervention in age-related retinal diseases, (e.g., AMD) and broadly from blood-CNS barriers in other neurodegenerative diseases.

## Introduction

The retinal pigmented epithelium (RPE) is a polarized cell monolayer in the eye that is situated between the photoreceptors and the systemic circulation of the choroid. The RPE is the initial site of pathological changes in age-related macular degeneration (AMD), which is the leading cause of blindness in people 65 years of age or older in developed countries (1). RPE are highly specialized cells that carry out crucial functions in the eye, including daily phagocytosis of outer segments shed from rod and cone photoreceptors, processing and transport of nutrients, and recycling of visual pigments (2). In addition, the RPE forms the outer blood-retinal barrier (oBRB) in the eye and its polarity is responsible for the directional secretion of proteins, lipoprotein particles and lipid bilayer-enclosed extracellular vesicles (EVs), including exosomes. Such polarity dictates directed interactions between the systemic circulation (basolateral) and the retina (apical). The role of this extensive endocytic trafficking, including the formation and release of a range of EVs, in AMD and other retinal diseases has not been thoroughly investigated to date (3).

Exosomes are cell-derived, bilayer-enclosed, nanovesicles (diameter (ø) = 30-150 nm) that are secreted in a controlled manner from most cell types. They make up the smallest subpopulation of the wide range of EVs released from most cells. It has become increasingly clear in recent years that exosomes have specialized functions and play a key role in, among other things, intercellular signaling, and cellular waste management (4). The results from a number of studies suggest that exosomes are not secreted merely as a degradation route for redundant molecules (5); rather they are equipped to withstand lysis by the complement system to carry out extracellular functions (6). Exosomes are formed inside specialized endosomes called multivesicular endosomes (MVE) and are released into the extracellular milieu upon MVE fusion with the plasma membrane. Their biogenesis and extracellular release is distinct from other EVs such as larger microvesicles that bud directly from the plasma membrane (7). Exosomes and microvesicles are functionally distinct in many respects (7). The role of exosomes and other EVs in the healthy and diseased eye has only recently begun to undergo rigorous study (reviewed in (3)). We and others have shown that highly polarized cells such as epithelia release exosomes and EVs in a directional manner with different cargoes detected in apical versus basolateral exosomes (8–17). However, there is a paucity of such studies to date using a global approach to characterize the changes to exosome protein content in response to common environmental stressors, such as chronic oxidative stress.

Oxidative stress has been implicated in RPE dysfunction that underlies development and progression of AMD (18–23). This dysfunction manifests in several RPE cellular processes including mitochondrial dysfunction (24), lipid metabolism perturbation (23), and disrupted autophagic clearance (25, 26). To date, most studies of experimental oxidative stress in RPE have only utilized acute and severe oxidative stress of 24-, 48-, but in rare cases, 72-hour treatments (27, 28) have been utilized. Longer experiments using milder oxidative stressors that more closely mimic early clinical stages of AMD, have rarely been conducted (25, 29). Cells under stress are known to increase the release of membranous vesicles including exosomes (30), and this has also been shown in RPE cells (31). There have been several recent studies analyzing the release of EVs from RPE cells under various types of stress, including acute and aggressive oxidative stress (29, 32, 33). However, to more closely mimic the early stages of RPE dysfunction in early dry AMD, we optimized conditions to induce a chronic subtoxic level of oxidative stress with minimal to no cell death. This was achieved using our previously validated primary porcine culture model of polarized RPE cell monolayers grown on semi-permeable membrane supports (12, 34, 35) and subjected to up to 4 weeks of low concentrations of hydrogen peroxide (H_2_O_2_). These chronically stressed primary RPE cultures on permeable transwell supports allow collection of separate EV populations secreted from the apical *and* basolateral side, which is essential for examining these epithelial monolayer responses in a more relevant physiological context.

Previous studies of secreted exosome and small EV preparations have been limited due to reliance on traditional mass spectrometric analysis of preparations of varying enrichment and purity, which are inherently heterogeneous mixtures, and thus reduce identification of low abundance proteins that are nonetheless specific for exosomes and small EVs. To address these limitations, we used density gradient flotation ultracentrifugation to achieve a higher degree of purity (36). In addition, to increase yield but maintain purity, we incorporated a novel step concentrating exosomes in a liquid cushion prior to density gradient ultracentrifugation.

Multiple lines of evidence indicate that one of the major culprits observed in RPE dysfunction is dysregulation in the endosomal pathway, which is thought to contribute to formation of ‘drusen’ (protein- and lipid-rich extracellular deposits associated with AMD) between the basal lamina of the RPE and the pentalaminar collagen- and elastin-rich Bruch’s membrane (BrM) (1). At present, the exact mechanisms for drusen formation are unknown. Since RPE-released exosomes and other (EVs) are essential parts of the endosomal pathway via their formation in MVEs that also provide trafficking of intracellular cargo (37), we undertook this study of exosomes released from chronically oxidatively stressed RPE cells, and the role they play in ECM changes that underlie early and late stages of AMD.

In the current study, we show for the first time that chronic subtoxic oxidative stress in primary RPE cultures induces changes in the protein cargo of exosomes released on the basal side reflecting desmosome and hemidesmosome shedding. Desmosomes are specialized and highly ordered membrane domains that mediate cell-cell contact and strong adhesion on the lateral cell surface (38). Hemidesmosomes on the other hand, are multiprotein complexes that facilitate the stable adhesion of epithelial cells via their basal surface to the underlying basement membrane (39). We also show that release of these exosomes correlates with ECM changes that can be prevented by inhibition of exosome release. These findings provide a novel avenue for therapeutic intervention and access to early biomarkers of cellular dysfunction in aging-related retinal diseases, in particular AMD.

## Materials and Methods

We have submitted all relevant data of our experiments to the EV-TRACK knowledgebase (EV-TRACK ID: EV230370) (40).

### Antibodies and reagents

Calcium and magnesium free PBS (PBS; #10010-023), and Hoechst 33258 (#H3569) were from Invitrogen (Waltham, MA). Triton X-100 (#T8787) was obtained from Sigma-Aldrich (St. Louis, MO). Antibodies used in the current study were as follows: Mouse anti-RPE65 (#NB100-355) [clone 401.8B11.3D9]; Novus Biologicals, Centennial, CO), mouse anti-DSG1 (#610273, BD Transduction, Franklin Lakes, NJ), mouse anti-Cytokeratin 10 (#MA5-13705 [clone DE-K10]; ThermoFisher, Waltham, MA), rabbit anti-Syntenin-1 (#ab19903]; Abcam, Cambridge, MA), mouse anti-Annexin II (#610068, BD Transduction Laboratories, Franklin Lakes, NJ), rat anti-Integrin Beta 1 (AIIB2 clone, Developmental Studies Hybridoma Bank, Iowa City, IA), mouse anti-Occludin (#66378-1-Ig, Proteintech, Rosemont, IL), rabbit anti-Calreticulin (#12238 [clone D3E6]; Cell Signaling Technologies, Danvers, MA), HRP-conjugated donkey-anti-rat IgG (#712-035-153, Jackson ImmunoResearch Laboratories, West Grove, PA), HRP-conjugated donkey-anti-mouse IgG (#715-035-150, Jackson ImmunoResearch), and HRP-conjugated donkey-anti-rabbit IgG (#711-035-152, Jackson ImmunoResearch).

### Polarized porcine RPE cell culture

Primary cultures of porcine RPE cells were prepared as described previously (12) with minor modifications. Briefly, porcine eyes were trimmed of excess tissue and anterior segments (including the entire lens and vitreous) were removed with a scalpel at the *ora serrata*. Eyecups (posterior poles) were placed into individual wells of 6-well tissue culture cluster plates (Corning #3516). Eyecups were filled with 2ml of PBS containing 1 mM EDTA (pre-warmed to 37°C) and incubated in a 37°C 5% CO_2_ incubator for 30 min to loosen the retina. After removal of the retina, eyecups were filled with 2ml of 0.25% trypsin-100 mM EDTA solution (Gibco #25200-056) and placed in an incubator at 37°C for 30 min. RPE cells were recovered by repeated aspiration followed by a low-speed centrifugation (5 min at 300 *g*). RPE cells were seeded at 60% confluence on 75 cm^2^ cell culture flasks (Corning #430641) and allowed to grow to >90% confluence before being trypsinized for seeding onto cell culture inserts. Thus, cells were seeded at passage p1 onto Laminin/Entactin (Corning #354259) coated 24 mm cell culture inserts with pore size of 0.4 µm (Corning Transwell™, #3450). Cells were seeded at high density onto inserts (300,000 RPE cells/cm^2^). Under these high-density seeding conditions, 100% confluence is achieved immediately upon seeding. RPE cells on inserts were maintained in DMEM with glucose and sodium pyruvate (Gibco #11995-065) supplemented with 1% (v/v) heat-inactivated FBS (Mediatech #35-010-CV), 100 units/ml penicillin, 100 µg/ml streptomycin, 2 mM L-glutamine (Sigma #G6784), MEM non-essential amino acids (Gibco #11140050), 0.25 µg/ml Amphotericin B (Gibco #15290-018), and 10 µg/ml Ciprofloxacin (Corning #61-277-RF). This medium will be referred to as *complete pig RPE (pRPE) medium* with FBS. High levels of pigmentation were achieved after as little as 1-2 weeks on cell culture inserts.

### Human RPE cell cultures from induced pluripotent stem cells (iPSC)

For the generation of iPSC-RPE cells with a known genetic risk for AMD, patients were recruited at the University Eye Clinic Regensburg after thorough clinical examination by an experienced ophthalmologist, the tests including fundoscopy and color fundus imaging (FF450plus fundus camera, Zeiss, Oberkochen, Germany), macular optical coherence tomography, fundus autofluorescence, as well as fluorescein angiography (all done with a Spectralis device, Heidelberg Engineering, Heidelberg, Germany). DNA samples were extracted from blood samples or skin biopsies and genotyped for 13 selected AMD-associated SNPs at 8 different loci known to be highly correlated with AMD risk by Sanger sequencing or restriction fragment length polymorphisms (RFLP) (41). Genetic risk scores were calculated using the model reported by Grassmann *et al.* (42) and samples from four high-risk (group 5) and four low-risk (group 1) individuals were included in the study. The culture and differentiation of human iPSC (hiPSC) from human donor material were approved by the Ethics Review Board of the University of Regensburg, Germany (reference no. 12-101-0241). Fibroblasts or peripheral blood mononuclear cells (PBMCs) from the selected donors were reprogrammed to iPSCs and subsequently differentiated into iPSC-RPE as described. Detailed information on the cell lines is given in (43). iPSC-RPE cells were thawed, seeded on 6-well plates coated with Matrigel® GFR (Corning Inc., Corning, NY), passaged once after 2 weeks and cultivated for another two weeks before seeding the cells onto transwell filters (ThinCert® Cell Culture Inserts by Greiner Bio-One, Kremsmünster, Austria). Conditioned apical and basal media were collected from cells which had been matured on transwell inserts for at least 6 and up to 9 weeks. Media was changed every 72 hours at the latest. After collection, media were centrifuged at 2,000 x *g* for 10min and then stored at -80°C until use for exosome isolation.

### Induction of oxidative stress

Oxidative stress conditions were induced by adding H_2_O_2_ daily to both the apical and basal compartments of fully differentiated RPE transwell cultures. The concentration was chosen from a range of concentrations tested from 50 to 500 µM, to achieve subtoxic oxidative stress conditions mimicking early stages of RPE dysfunction without catastrophic loss of barrier integrity (occurs at TER below 100 Ω•cm^2^; (44)) and outright RPE dysfunction and cell death within the 4-week experimental treatment duration. The identified concentration was 0.2 mM H_2_O_2_.

### Transepithelial electrical resistance (TER)

TER) is a reliable assay for the assessment of RPE barrier function and is inversely proportional to the paracellular permeability of cultured RPE cells (45–48). It was measured by means of a volt-ohm meter (EVOM) equipped with a 24 mm EndOhm chamber (both from World Precision Instruments, Sarasota, FL). Resistance values for each condition were determined from a minimum of three individual cultures and corrected for the inherent cell culture insert resistance within five minutes after removing the plates from the incubator. All values represent the mean ± S.E.M.

### LDH cytotoxicity assay

Twenty microliter aliquots of apical media from each well in a Control, and an H_2_O_2_-treated 6-well transwell plate, were collected before first treatment (day 0), and weekly thereafter for the 4-week experiment. Media aliquots were collected from media that had been conditioned with the cells for 48 hours, in all cases. Aliquots were frozen at -80°C until analysis. Amounts of LDH in conditioned media were assessed by using an ultrasensitive bioluminescent plate-based assay (LDH-Glo™ Cytotoxicity Assay, Promega, Madison, WI). We followed the manufacturer’s instructions in their entirety and used a Molecular Devices SpectraMax M5 (San Jose, CA) multimodal microplate reader for luminescence signal detection.

### Conditioned media for EV isolation

For generation of conditioned media for EV isolation, cell media was exchanged with media supplemented with 2% (v/v) EV-depleted FBS. To avoid contamination with FBS-derived EVs from the complete pRPE medium, cells were cultured for one day in the EV collection media after which it was discarded, and fresh collection media was added. EV-depleted FBS was prepared as described previously (49). Briefly, 20% (v/v) FBS was centrifuged in a Beckman Optima XE-90 ultracentrifuge using an SW 28 Ti rotor at 100,000 *g*_avg_ for 18 hrs at 4°C. The supernatant was carefully collected without disturbing the loose pellet and sterile-filtered through a 0.22 µm PVDF filter bottle (Millipore), aliquoted and frozen at -20°C until used.

For EV collection from cell culture inserts: For each treatment condition, conditioned media from two 6-well cluster plates with 24mm permeable inserts were collected every other day (every 48 hours) for four weeks. The volumes used were 1.5ml in the upper (apical) chamber and 2.6ml in the lower (basal) chamber for each insert. For an average exosome preparation, 100-200ml of apical and 200-400ml of basal conditioned media was used as starting material.

### EV isolation

Two experimental protocols were used to isolate EVs:

#### EV isolation by differential centrifugation

EVs were isolated using a modification of a well-established differential centrifugation protocol as previously described (49, 50). Briefly, conditioned medium was centrifuged at 2,000 *g* for 10 min to remove cell debris and the resulting supernatant was collected and kept at -80°C until the next steps of the isolation protocol. Cleared conditioned media was centrifuged at 10,000 *g* for 30 min and the resulting supernatant was transferred to a new tube and centrifuged at 100,000 *g*_avg_ for 90 min. The resulting supernatant was discarded, and the EV pellet was resuspended in a lysis buffer (2% SDS, 100 mM Tris-HCl [pH 6.8]) if the sample was prepared for immunoblotting analyses or resuspended in PBS if prepared for NTA analysis. Centrifugations at 10,000 *g* (*k*-factor = 2547.2) and 100,000 *g* (*k*-factor = 254.7) were done at +4°C using polyallomer tubes (Beckman Coulter Inc., Indianapolis, IN; #326823) in an SW 28 Ti rotor in a Beckman Optima XE-90 ultracentrifuge (Beckman Coulter).

#### EV isolation by cushioned iodixanol buoyant density gradient centrifugation

Conditioned media previously cleared at 2,000 *g* was concentrated using Centricon Plus-70 centrifugal filter devices with 100 kDa NMWL cutoff (Millipore, #UFC710008), or Corning Spin-X 20ml centrifugal filter devices with 100 kDa NMWL cutoff (SigmaAldrich, #CLS431491). We modified a previously published protocol from (51, 52) to provide a gentler EV isolation with improved yield, see schematic of workflow in **Fig. 1**. Briefly, the concentrated media was centrifuged at 10,000 *g* for 30 min as described in the previous paragraph, and the resulting supernatant was recovered and made up to 36ml with PBS if needed. The supernatant was carefully placed in a polyallomer tube onto a 2ml cushion of 60% OptiPrep (Sigma-Aldrich #D1556) and centrifuged at 100,000 *g_avg_* in an SW 28 Ti rotor for 180 min. The resulting 2ml cushion and 1ml of interface was collected from the bottom using a 4-inch blunt 18-gauge metal hub Luer lock needle (Hamilton Company, #7748-04) with a 5ml syringe (BD#309646) extending from the open top. The collected 3ml was used as the bottom 40% OptiPrep fraction in the subsequent density gradient. A discontinuous gradient of iodixanol solutions was then prepared by carefully overlaying the bottom fraction with 3ml of 20%, 10%, and 5% solutions of iodixanol buffered with [0.25M sucrose, 10 mM Tris-HCl (pH 7.5)], respectively. The gradient tubes (UltraClear™; Beckman Coulter #344059) were subjected to centrifugation at 200,000 *g_avg_* (SW 41 Ti rotor; 40,000 rpm) for 4 hrs at +8°C. One ml fractions were collected manually from the top of the self-generated gradient and weighed to determine density. One ml fractions were diluted 12-fold with PBS and subjected to centrifugation at 100,000 *g_avg_* for 90min in the SW 41 Ti rotor. Pellets were resuspended in 50-100μl 2% SDS, 100 mM Tris-HCl [pH 6.8] and stored at -80°C until use. Total protein content in EV preparations were determined with the Pierce 660nm protein assay (ThermoFisher Scientific #22660) using a NanoDrop 2000 spectrophotometer (ThermoFisher Scientific #ND-2000).

**Figure 1.**
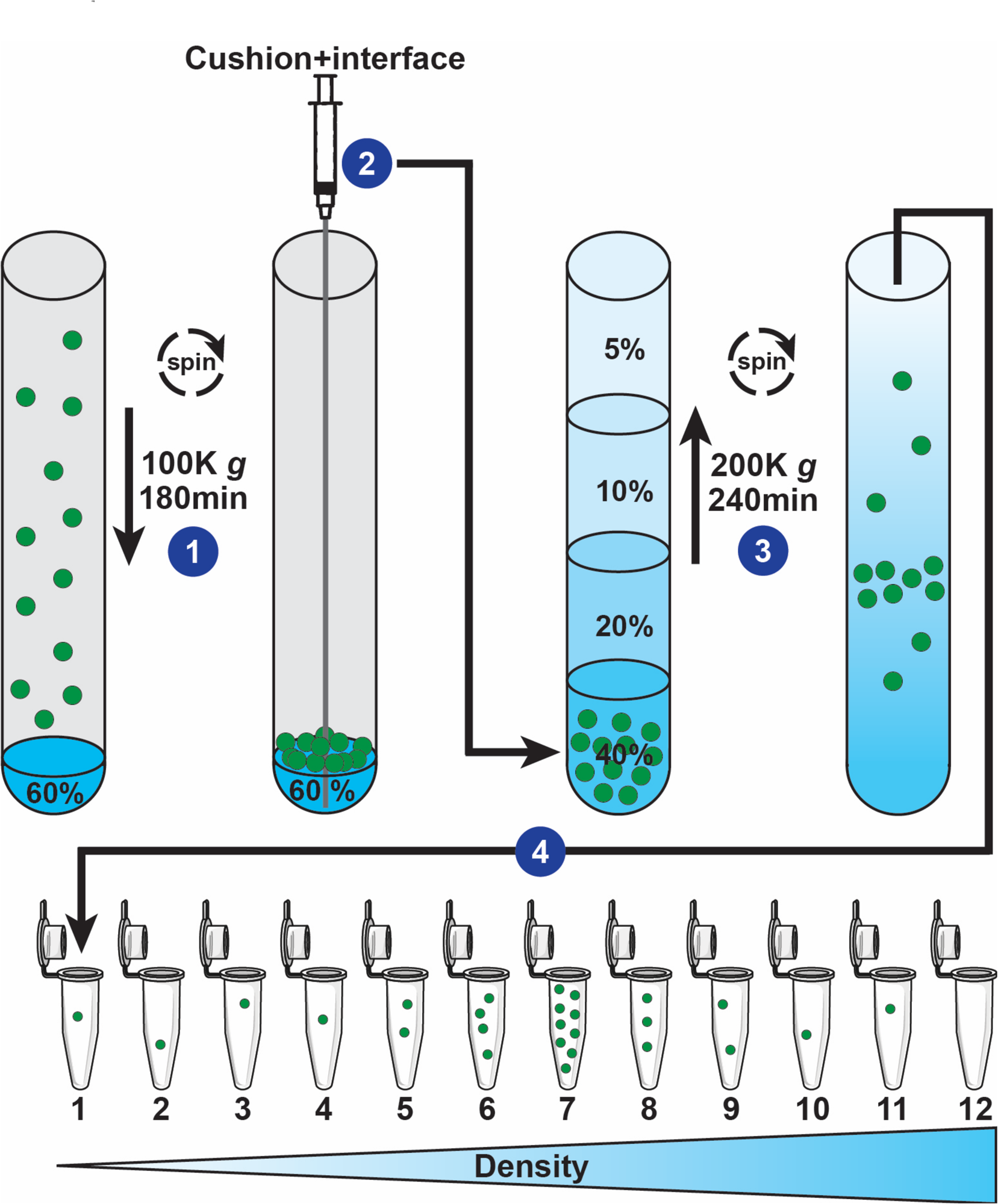
Schematic of a cushioned flotation density gradient ultracentrifugation (C-DGUC) method for isolation of small EVs (sEV) including exosomes. (**1**) EV-containing fluid that has already undergone sequential centrifugations of 2,000 *g* and 10,000 *g* to remove cell debris and large EVs, is placed on top of a cushion of 60% Iodixanol (OptiPrep^TM^) and sEV are pelleted by ultracentrifugation. (**2**) 60% Iodixanol cushion and interface are collected from the bottom using a 4-inch blunt needle extending from the open top. The collected sEV 40% Iodixanol sample is placed on the bottom of a tube and (**3**) carefully overlaid with three Iodixanol step fractions of decreasing concentrations (20%, 10%, and 5%). Flotation of EVs to their intrinsic density is achieved by ultracentrifugation of the self-forming gradient. (**4**) Twelve fractions of equal volume are collected from the top of the gradient, and weighed or analyzed by spectrophotometry for Iodixanol content, to determine density and identify location of exosomes (ϱ=1.07-1.11 g/ml). Individual fractions are processed for downstream analyses.

### Immunoblotting

Western blot analysis was performed as previously described (53). Briefly, cell lysate, ECM, crude EV and purified iodixanol gradient fraction samples were run reduced or unreduced, on 10% or 4-12% gradient Bis-Tris Criterion XT gels in MOPS buffer, transferred to PVDF using a BioRad Trans-Blot Turbo Semi-Dry transfer apparatus, and then probed with indicated antibodies. Anti-Syntenin-1, anti-RPE65, and anti-Calreticulin were used at 1:1,000 dilutions. Anti-Annexin II, anti-Cytokeratin 10, and anti-Desmoglein 1 were used at dilutions at 1:500. Anti-Occludin was used at 1:3,000 dilution. Anti-Integrin Beta 1 was used at 1:88 dilution. Subsequent incubation with horseradish peroxidase-conjugated secondary antibodies at 1:40,000 dilution was followed by detection with SuperSignal West Pico Plus (ThermoFisher Scientific #34580) or Immobilon ECL Ultra Western HRP Substrate (Sigma #WBULS0500). ECL signals and total protein loading amounts were measured with a Bio-Rad ChemiDoc MP imaging system (Bio-Rad Laboratories Inc., Hercules, CA). The acquired images were analyzed with Bio-Rad ImageLab software version 6.1 (Bio-Rad Laboratories).

### EV sample preparation for proteomics

EV samples in lysis buffer (2% SDS, 100 mM Tris-HCl [pH 6.8]) were prepared for proteomic analysis and digested with trypsin using a recently reported paramagnetic bead-based protocol (54) that is described in detail in (12). Three porcine RPE exosome preparations (three biological repeats) of basolateral origin and two of apical origin, were analyzed by mass spectrometry. Two hiPSC-RPE exosome preparations of basolateral origin from each of four high genetic AMD-risk lines and four low genetic AMD-risk lines, were analyzed by mass spectrometry. For all discussions of proteomic data, the term ‘set(s)’ is used throughout the current study to describe individual mass spectrometry datasets from biological replicate EV preparations.

### EV sample preparation and LC-MS/MS analysis

Proteins were cleaved with the trypsin/endoproteinase LysC mixture (Promega, V5072) using the paramagnetic bead-based method (54). Each digest was dissolved in 12 μl of 1/2/97% (by volume) of the trifluoroacetic acid/acetonitrile/water solution, and 3 μl were injected into a 5 μm, 180 μm × 20 mm Symmetry C18 trap column (Waters) in 1% acetonitrile in water for 3 min at 5 μl/min. The analytical separation was next performed using an HSS T3 1.8 μm, 75 μm × 200 mm column (Waters) over 90 min at a flow rate of 0.3 μl/min at 55°C. The 5-30% mobile phase B gradient was used, where phase A was 0.1% formic acid in water and phase B 0.1% formic acid in acetonitrile. Peptides separated by LC were introduced into the Q Exactive HF Orbitrap mass spectrometer (ThermoFisher Scientific) using positive electrospray ionization at 2000 V and capillary temperature of 275°C. Data collection was performed in the data-dependent acquisition (DDA) mode with 120,000 resolution (at m/z 200) for MS1 precursor measurements. The MS1 analysis utilized a scan from 375-1450 m/z with a target AGC value of 1.0 x 10^6^ ions, the RF lens set at 30%, and a maximum injection time of 50 ms. Advanced peak detection and internal calibration (EIC) were enabled during data acquisition. Peptides were selected for MS/MS using charge state filtering (2–5), monoisotopic peak detection and a dynamic exclusion time of 25 sec with a mass tolerance of 10 ppm. MS/MS was performed using higher-energy C-trap dissociation (HCD) with a collision energy of 30±5% with detection in the ion trap using a rapid scanning rate, automatic gain control target value of 5.0 x 10^4^ ions, maximum injection time of 150 ms, and ion injection for all available parallelizable time enabled.

### EV Protein identification and quantification

For label-free relative protein quantification, raw mass spectral data files (.raw) were imported into Progenesis QI for Proteomics 4.2 software (Nonlinear Dynamics) for duplicate runs alignment of each preparation and peak area calculations. Peptides were identified using Mascot version 2.6.2 (Matrix Science) for searching Sus scrofa UniProt July 2019 database. Mascot search parameters were: 10 ppm mass tolerance for precursor ions; 0.025 Da for fragment-ion mass tolerance; one missed cleavage by trypsin; fixed modification was carbamidomethylation of cysteine; variable modification was oxidized methionine. Only proteins identified with 2 or more peptides (protein confidence p<0.05 and false discovery rate <1%), were included in the protein quantification analysis. To account for variations in experimental conditions and amounts of protein material in individual LC-MS/MS runs, the integrated peak area for each identified peptide was corrected using the factors calculated by automatic Progenesis algorithm utilizing the total intensities for all peaks in each run. Values representing protein amounts were calculated based on a sum of ion intensities for all identified constituent non-conflicting peptides (55).

### Isolation of RPE ECM for immunoblotting analysis

Primary porcine RPE cells were grown to confluence in 6-well Transwell inserts, media was removed, and wells washed twice with 1ml PBS per well. Each well was placed on individual small petri dishes to avoid liquid leaking through. One ml of [20 mM NH_4_OH, 0.5% Triton X-100] was added to each well to lyse cells and incubated for 15 min at room temperature (RT). Wells were rocked occasionally to ensure entire area of wells were lysed. Solution was removed, and wells washed twice carefully with 1 ml ddH_2_O per well. 200µl of 2x GLB + 8M Urea lysis buffer was added to each well on petri dishes. Wells were incubated at RT for 10min. Wells were again rocked occasionally to ensure entire area of wells were solubilized. ECM solution was scraped with cell scraper and collected in 1.5ml tubes. Tubes were vortexed vigorously, then insoluble debris was cleared at 10,000 *g* for 15min at +4°C. Supernatant was frozen in a fresh tube at -80°C.

### Isolation and preparation of RPE ECM for mass spectrometry analysis

Cell culture flasks were decellularized using the method of Vlodavsky *et al.* (56). Briefly, human (fetal human RPE were provided and isolated by A. Maminishkis as previously described in (57)) and porcine primary RPE cells were grown to confluence in T-75 flasks and washed once with PBS. Four ml of [20 mM NH_4_OH, 0.5% Triton X-100] was added to lyse cells and incubated for 30 min at RT. Twenty ml of de-ionized H_2_O was added to flask, with rocking. Liquids were removed by vacuum aspiration with a glass Pasteur pipette. The insoluble ECM layer was washed with 20 ml de-ionized H_2_O and repeated 4 more times for a total 5 washes to ensure the complete removal of NH_4_OH and Triton X-100. One ml [8M Urea, 50 mM Tris-HCl (pH 8.0)] was added and incubated with gentle shaking for 5min to denature ECM protein content. To reduce disulfide bonds, Dithiothreitol (DTT) was added to a final concentration of 5 mM and incubated for 30min at 37°C. The reduced thiol group was alkylated by addition of Iodoacetamide (IAA) to a final concentration of 15 mM and incubated in the dark at RT for 30min. Mass spectrometry grade Trypsin or Trypsin/Lys-C Mix (Promega #V5111 or V5071) was added at to a final concentration of 1µg/ml and incubated for 30 min at 37°C in the flask. The reaction was then diluted 8-fold with 50 mM Tris-HCl (pH 8.0) to reduce Urea concentration to 1M which allows sufficient Trypsin activity for efficient cleavage. ECM was collected with a cell scraper together with the 8ml solution and incubated overnight at 37°C. The reaction mixture was centrifuged at 10,000 *g* for 15 min at RT to pellet insoluble material. The supernatant was collected and Trifluoroacetic acid (TFA) was added to 0.2% (v/v) to terminate trypsinization. Peptide samples were cleaned using C18 Silica tip columns (The Nest Group Inc., Southborough, MA; #SEM SS18V). C18 columns were wetted with 200µl 40% ACN, 40% MeOH, followed by equilibration twice with 200µl 2% ACN, 0.2% TFA. The 8ml sample was then added in 200µl portions, 3-4 drops at a time, incubated for 1min, then repeated until entire sample had been applied. Columns were then washed with 3 portions of 200µl 2% ACN, 0.2% TFA. Finally, the cleaned peptides were eluted with 200µl 50% ACN, 0.2% TFA and vacuum-dried using an Eppendorf Vacufuge (#22822128) with 45°C heat until dry for 30min.

Dried samples were resuspended by adding 2µl (20µg) of a mixture of hydrophilic and hydrophobic paramagnetic beads as previously described (54). Briefly, the mixture was acidified by adding formic acid to 0.25% (v/v) and mixed by pipetting to resuspend dried sample on tube walls. 195µl of acetonitrile was added to a final concentration of >95% and mixed carefully by pipetting up and down. Samples were incubated for 10 min, briefly centrifuged to pellet beads, and tubes were placed on magnetic rack. Following a 2 min incubation, the solution was carefully withdrawn and discarded. Beads were rinsed while on magnet with 180µl of acetonitrile by pipetting up and down multiple times. Following a 2 min incubation, the solution was again carefully withdrawn and discarded, and tubes were briefly air-dried. Peptides were eluted by adding 20µl 2% DMSO, 0.2% formic acid (FA) and pipetting beads up and down. Following a 5 min incubation, tube was placed on magnetic rack and the eluate was carefully collected. The elution procedure was then repeated with the addition of 20µl 0.2% FA and carefully collected and combined with the first eluate. Eluate was vacuum-dried, resuspended in 20µl ddH_2_O and peptide concentration was determined by A280 quantitation using a NanoDrop 2000 spectrophotometer.

### Nanoparticle tracking analysis

Nanoparticle tracking analysis (NTA) was performed using a ZetaView PMX110 Instrument (ParticleMetrix, Inning am Ammersee, Germany) equipped with a 405 nm laser and integrated automated fluidics. Three cycles of recordings at the ‘high’ setting (2-second videos) were collected of each sample with camera sensitivity set at 92, shutter at 70, minimum brightness at 15, minimum size at 10 and maximum size at 5,000. Temperature was set at 26°C and monitored throughout the measurements. Videos recorded for each sample were analyzed with ZetaView software version 8.04.02 SP3 to determine the concentration and size of measured particles with corresponding standard error. For analysis, auto settings were used for blur, minimum track length and minimum expected particle size. The ZetaView system was calibrated with 102nm polystyrene microbeads (ThermoFisher Scientific #3100A) prior to every analysis session. PBS (Gibco #10010-023) was used as diluent, and to avoid contaminating particles, a fresh bottle was opened for each analysis session. At least three separate EV preparations for each condition were analyzed.

### Statistical analysis

TER measurements, concentrations of small EVs released from RPE, modal and mean EV sizes, and immunoblot densitometry quantitation were tested for statistical significance using a two-sample two-tailed Student’s t-test assuming unequal variance; two-way or one-way ANOVAs with corrections for multiple corrections were used when appropriate. P-values < 0.05 were considered statistically significant. All statistical analyses were performed using Prism Software (GraphPad Prism Version 9.4.1 (458) for MacOS, San Diego, CA).

#### Additional apical exosome proteome statistical analysis

Raw peptide abundances collected from mass spectrometry of H_2_O_2_-treated (n=2) and untreated (n=2) apical exosomes were analyzed using DESeq2 (58). Technical replicates (n=2) for each sample were combined using collapseReplicates-DESeq2 and a median of ratios normalization was applied to generate relative protein abundances. Differences in protein abundance by H_2_O_2_-treatment condition were detected using a likelihood-ratio test. P-values were adjusted using an FDR cutoff of 0.1.

## Results

### Chronic subtoxic oxidative stress conditions do not affect tight junction integrity or canonical RPE protein expression

We developed a chronic subtoxic oxidative stress model of RPE by treating our highly differentiated polarized porcine RPE (pRPE) monolayer transwell cultures (12) daily with 0.2 mM H_2_O_2_ and collecting conditioned every 48 hours from Control and Treated, for 4 weeks. The concentration was chosen from a range of concentrations of H_2_O_2_ as the one that achieved subtoxic oxidative stress conditions, mimicking early stages of RPE dysfunction, without loss of barrier integrity or outright RPE dysfunction and cell death. Fully differentiated RPE cultures form tight junctions which give rise to an epithelial fluid and ion barrier which can be assessed by measuring the trans-epithelial electrical resistance (TER). Stressed or diseased RPE monolayers lose some barrier integrity that result in decreased TER (44). We monitored TER in the stressed RPE cultures over the 4 weeks to confirm that epithelial barrier function was maintained above loss of barrier integrity, which occurs below 100 Ω•cm^2^, for the duration of the experiment (**Fig. 2A**). At the light microscopy level there was no detectable disruption of RPE cell morphology following 4 weeks of 0.2 mM H_2_O_2_ exposure (**Supp. Fig. S1A**), and this was further confirmed using a cell cytotoxicity assay measuring the release of Lactate Dehydrogenase (LDH (**Supp. Fig. S1B**).

**Figure 2.**
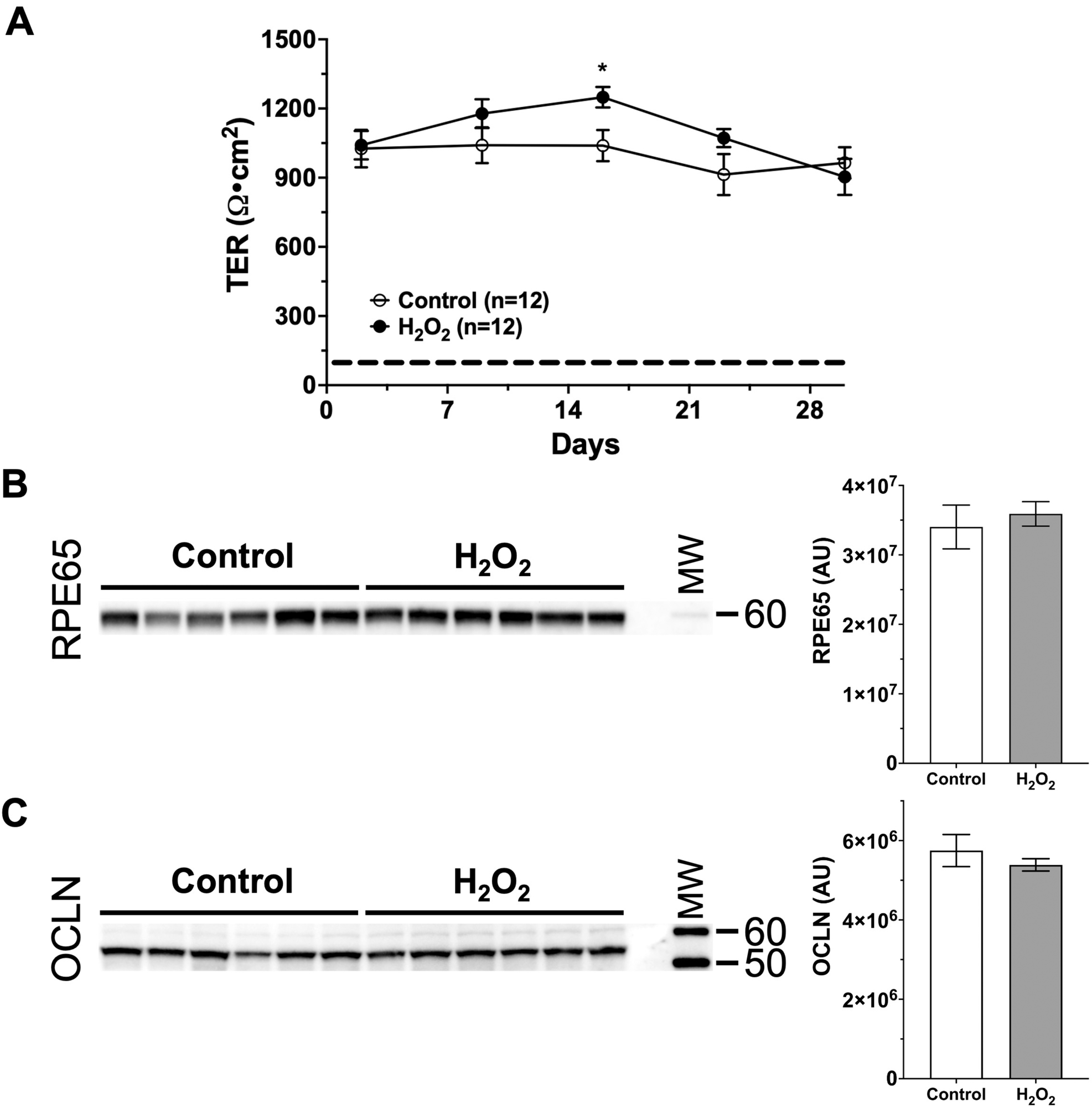
Chronic subtoxic oxidative stress conditions do not affect tight junction integrity or canonical RPE function. (**A**) To assess the integrity of pRPE monolayers, the transepithelial electrical resistance (TER) was measured weekly during the 4-week experiment. Throughout the experiment TER in H_2_O_2_-treated cultures remained at levels similar to control cultures and above the level indicating intact barrier integrity (dashed line). Values plotted are mean ± SEM, twelve replicates per data point. Data were analyzed by multiple unpaired t tests with Welch correction, assuming unequal variance. * = p < 0.05. (**B-C**) Ten micrograms of total protein in lysates (because oxidative stress affects most housekeeping proteins, they can’t be used as loading controls, thus matched loading of total protein is the preferred method) of Control and H_2_O_2_-treated RPE transwell cell cultures after 4-week experiments, were analyzed by immuno-blotting. Each lane contains a lysate from an individual well in 6-well transwell plates. (**B**) RPE65, which is essential for normal RPE function, did not change under these oxidative stress conditions. (Right panel) Quantitation of chemiluminescent immuno-detection signal. (**C**) Occludin (OCLN), which is a tight junction protein essential for RPE barrier integrity, did not change under these oxidative stress conditions. (Right panels) Quantitation of chemiluminescent immunodetection signal. Error bars are SEM. Unpaired Student’s t-test assuming unequal variance was used to assess statistical significance for treatment condition. MW = Molecular weight markers. Apparent molecular weights in kDa are indicated on the right-hand side. AU = Arbitrary units.

We hypothesized that the chronic stress induced by 4 weeks of 0.2 mM H_2_O_2_ treatment would have little to no detectable impact on normal cellular function since the cells remained morphologically normal with a normal TER throughout the oxidative stress treatment. To test this, we assayed RPE cell lysates isolated from pRPE with or without H_2_O_2_ for 4 weeks, by immunoblotting for changes to levels of two relevant markers: (1) RPE65, which is an enzyme essential for the visual cycle pathway in RPE cells (**Fig. 2B**), and (2) Occludin (OCLN), a tight junction protein essential for barrier integrity (**Fig. 2C**). The concentrations of both proteins remained unchanged under oxidative stress conditions compared to controls, indicating that these conditions were largely subtoxic.

### A chronic subtoxic oxidative stress RPE model increases basolateral exosome release and causes changes in the exosome proteome

EVs released basolaterally from control and H_2_O_2_-treated pRPE during weeks 1 & 2 and 3 & 4 were quantified by Nanoparticle Tracking Analysis (NTA). Basolateral EV release was significantly increased during weeks 3 & 4 of H_2_O_2_ treatment (**Fig. 3A**). No significant differences were observed in average or modal EV sizes under Control or H_2_O_2_ conditions (**Supp. Table S1**). NTA analyses of apical EVs showed a decreased release in response to H_2_O_2_ treatment (**Supp. Fig. S2**). Basolaterally released pRPE EVs isolated by cushioned Iodixanol density gradient ultracentrifugation (C-DGUC, **Fig. 1**) from fractions with densities corresponding to exosomes (**Supp. Fig. S3**), were shown by immunoblotting to contain the canonical exosome markers, Syntenin-1 and Annexin A2 (ANXA2) (**Fig. 3B-C**). The absence of the ER marker Calreticulin (CALR) in these EV preparations confirmed a lack of cellular contamination.

**Figure 3.**
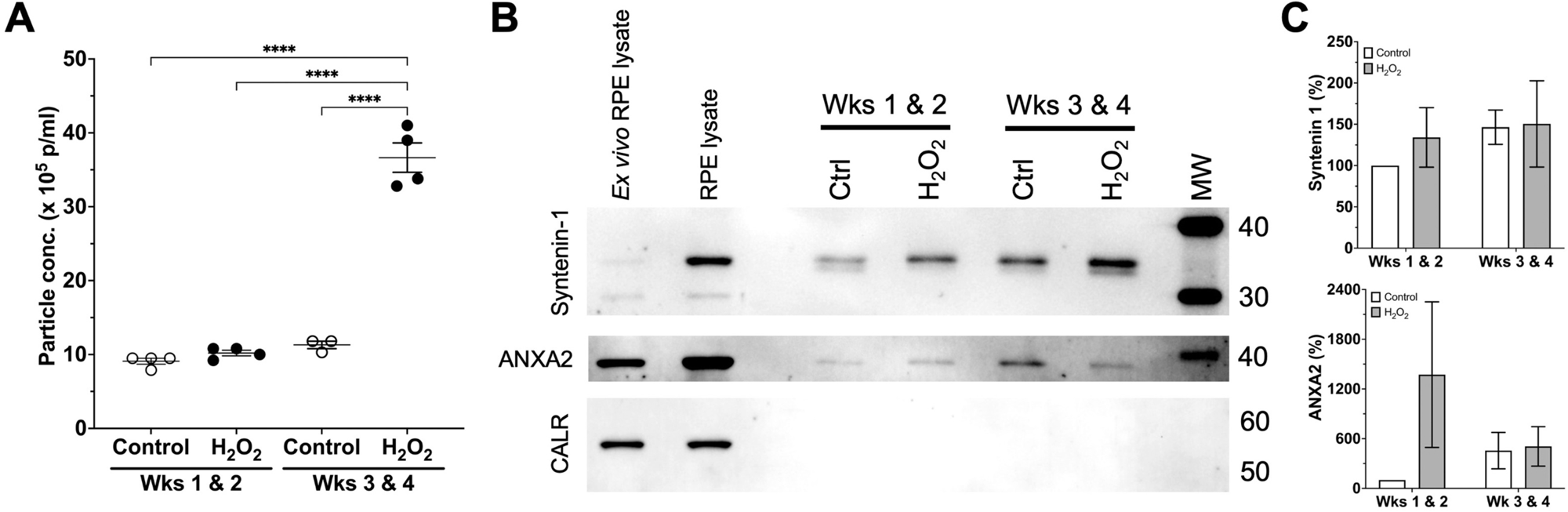
A chronic subtoxic oxidative stress RPE model increases basolateral exosome release and cause changes in canonical exosome protein markers. Fully differentiated and polarized pig RPE transwell cultures were treated or not for four weeks with 0.2 mM H_2_O_2_. (**A**) EVs released basolaterally from control and H_2_O_2_-treated pRPE during weeks 1 & 2 and 3 & 4 were quantified by NTA. EV release was significantly increased during weeks 3 & 4 of H_2_O_2_ treatment. Mean ± SEM is indicated. One-way ANOVA analysis for treatment condition followed by Tukey’s *post-hoc* multiple comparison analysis indicated as **** = p < 0.0001. (**B**) Representative immunoblots of basolateral pRPE EVs isolated by cushioned Iodixanol density gradient ultracentrifugation (C-DGUC). Equal volumes of EV lysates were used in these analyses. Exosome markers Syntenin-1 and Annexin A2 (ANXA2) demonstrates the presence of exosomes and sEV while the absence of the ER marker Calreticulin (CALR) in EV preparations indicates a lack of cellular contamination. Lysate of RPE cells freshly isolated from pig eyes (*Ex vivo* RPE lysate), and lysate of cultured pig RPE cells (RPE lysate) are indicated. Apparent molecular weights in kDa are indicated on the right-hand side. (**C**) Quantitation of chemiluminescent immunodetection signal density on immunoblots. All values are normalized to the ‘Control Wks 1 & 2’ EV preparation and expressed in percent. Quantitation of Syntenin-1 immunodetection is shown in the *upper panel*, and ANXA2 quantitation in the *lower panel*. Error bars are SEM.

We processed the entirety of these exosome preparations isolated by C-DGUC by an in-solution bead-based digestion protocol followed by electrospray-tandem mass spectrometric analysis. Basolateral exosomes from Control and H_2_O_2_-treated pRPE cultures were compared in a quantitative manner using the ProGenesis QI software suite. Proteins that were increased 2-fold or more in two sets (individual mass spectrometry datasets from biological replicate EV preparations) during weeks 3 & 4, and one set during weeks 1-4, are shown in **Table 1**. Proteins decreased two-fold or more are shown in **Supp. Table S3**. Proteomic analyses of basolateral exosomes released during weeks 1 & 2 were variable between sample sets and did not reveal any clear consensus among proteins increased or decreased 2-fold or more (data not shown).

**Table 1.**
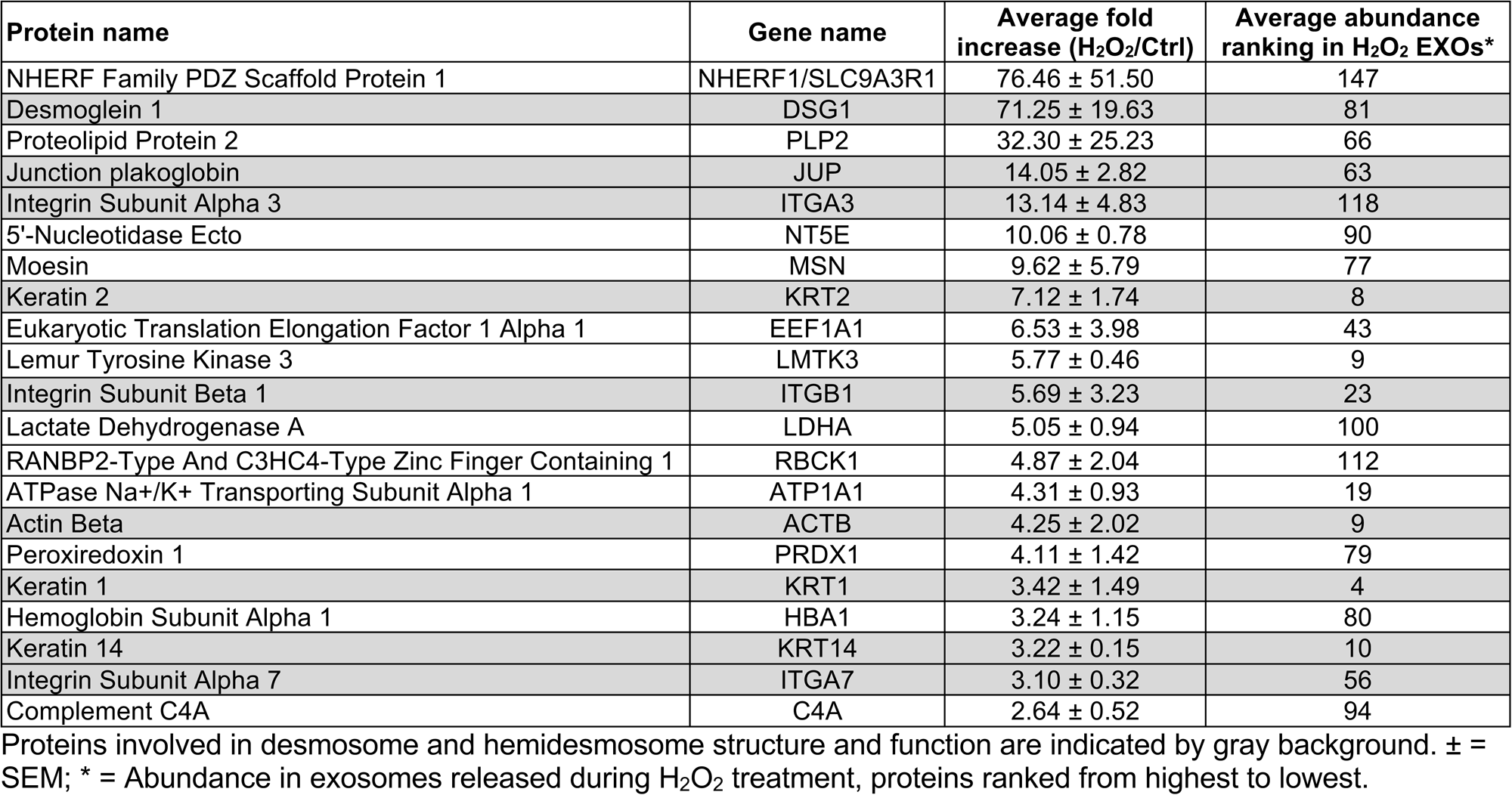
Proteins increased ≥2-fold in at least two of the three sets of <u>basolateral</u> exosomes released during weeks 3 & 4 from H_2_O_2_-treated RPE cultures compared to untreated. Sets #1 & #2 represent two biological replicates. Set #3 was generated from exosomes released during an entire 4-week experiment. Each set was generated from two technical mass spectrometric replicate runs.

Apical exosomes isolated in the same way as basolateral exosomes were also processed and analyzed the same way. The apical exosome proteins that were increased (**Table 2**) or decreased (**Supp. Table S4**) two-fold or more during weeks 3 & 4 of H_2_O_2_ treatment of the pRPE are shown.

**Table 2.**
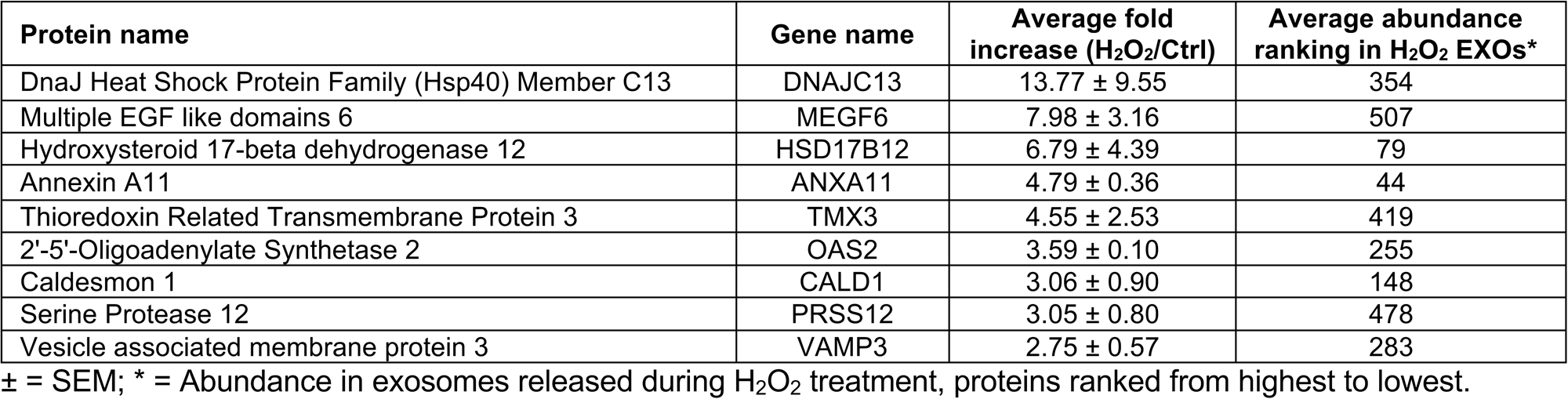
Proteins increased ≥2-fold in both sets of <u>apical</u> exosomes released during weeks 3 & 4 from H_2_O_2_-treated RPE cultures compared to untreated. Sets #1 & #2 represent two biological replicates. Each set was generated from two technical mass spectrometric replicate runs.

### Subtoxic oxidative stress induces basolateral exosome release of desmosomal and hemidesmosomal proteins

Analysis of the basolateral exosome proteome revealed that a set of proteins important for desmosomal and hemidesmosomal ECM-attachment structure and function were statistically significantly increased upon oxidative stress (see proteins highlighted by a gray background in **Table 1**). Functional enrichment analyses using the FunRich analysis tool (version 3.1.4) confirmed these observations and robustly identified the subcellular location ‘desmosome’ as statistically significantly increased in exosomes released under oxidative stress conditions (data not shown). Interestingly, enrichment of desmosomal or hemidesmosomal proteins were not seen in apical exosomes (**Table 2**), suggesting a highly polarized response to oxidative stress. An increase in the hemidesmosomal marker, Integrin beta 1 (ITGB1) in the protein cargo of basolaterally released EVs under oxidative stress was further validated by immunoblotting (**Fig. 4A**). It is also worth noting that ITGA3 and ITGB1, which form a preferred heterodimer (59, 60), were both increased in response to stress (**Table 1**).

**Figure 4.**
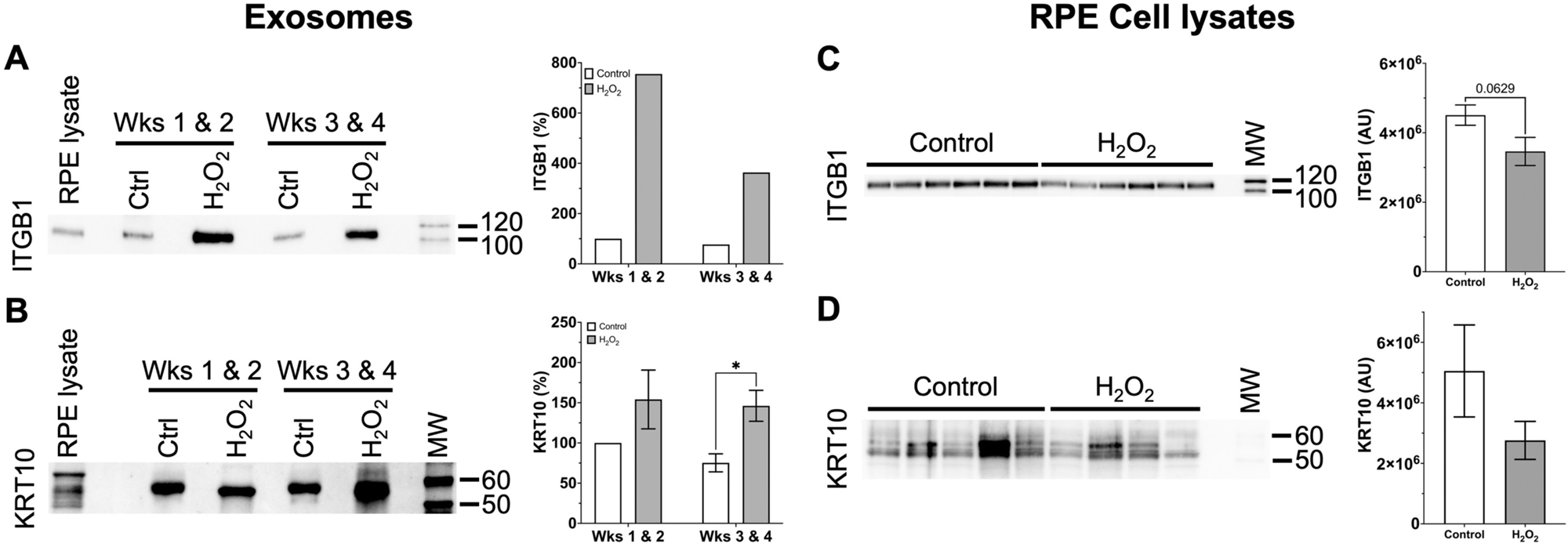
Immunoblotting validation of a subset of desmosomal proteins released via basolateral exosomes under subtoxic oxidative stressor conditions. (**A-B**) Basolaterally released exosomes were analyzed by immunoblotting for a subset of desmosomal proteins in control and H_2_O_2_-treated RPE cultures during the early (weeks 1 & 2) and late (weeks 3 & 4) parts of 4-week experiments. Representative immunoblots for (**A**) Integrin beta 1 (ITGB1), and (**B**) Keratin 10 (KRT10) show an increased abundance of these proteins in exosomes released from H_2_O_2_-treated RPE cultures. This correlated with findings from unbiased mass spectrometric analyses (see **Table 1**). Quantitation of chemiluminescent immunodetection signal on immunoblots are shown to the right in panels. All values are normalized to the ‘Control Wks 1 & 2’ EV preparation and expressed in percent. (**C-D**) Lysates of Control and H_2_O_2_-treated RPE transwell cell cultures after 4-week experiments were analyzed by immunoblotting. Each lane contains a lysate from an individual well in 6-well transwell plates. Representative blots for (**C**) ITGB1, and (**D**) KRT10 are shown. Quantitation of chemiluminescent immunodetection signal density on immunoblots was done as described in (**A-B**) and show a decrease in cells under oxidative stress conditions. Unpaired Student’s t-test assuming unequal variance was used to assess statistical significance for treatment condition. P-values below 0.1 are indicated. * = p < 0.05. AU = Arbitrary density units.

Immunoblotting for the desmosome and hemidesmosome-anchoring protein, KRT1 proved challenging due to a lack of high-quality antibodies against porcine KRT1. Therefore, given that KRT1 is known to form preferred heterodimers with KRT10, and these pairs play an essential role in desmosome structure integrity (61), we analyzed the quantity of KRT10 as a proxy for KRT1 in basal exosomes. KRT10 was only increased more than two-fold in one of three proteomic datasets (**Supp. Tables S8-10**), and was therefore not included in **Table 1**. However, KRT10 was increased in response to oxidative stress on immunoblots of protein lysates from three separate basal exosome preparations (**Fig. 4B**). This correlated with proteomic findings for KRT1 (**Table 1**).

Significantly, a similar subset of desmosomal proteins were increased in exosomes released basolaterally from untreated induced pluripotent stem cell (iPSC)-derived RPE from AMD patients with high genetic AMD risk compared to low-risk control lines (**Table 3**, **Supp. Tables S5, and S13-16**).

**Table 3.** Desmosome and hemidesmosome-associated proteins in <u>basolateral</u> exosomes increased ≥2-fold in at least two of four iPSC-derived RPE lines from donors with high genetic AMD risk compared to four iPSC-RPE lines from donors with low risk (42, 80). Data was generated from two technical mass spectrometric replicate runs.

| Protein name | Gene name | Average fold increase<br>(High-risk/Low-risk) | Average abundance<br>ranking in High-risk EXOs* |
| --- | --- | --- | --- |
| CD151 | CD151 | 144.69 ± 244.37 | 20 |
| Keratin 34 | KRT34 | 64.47 ± 43.70 | 120 |
| Junction Plakoglobin | JUP | 14.93 ± 11.96 | 126 |
| Keratin 5 | KRT5 | 6.27 ± 3.68 | 21 |
| Keratin 1 | KRT1 | 4.87 ± 2.88 | 3 |
| Keratin 16 | KRT16 | 4.84 ± 4.54 | 19 |
| Keratin 2 | KRT2 | 3.75 ± 1.56 | 9 |
| Desmoplakin | DSP | 3.47 ± 1.22 | 121 |
| Keratin 81 | KRT81 | 3.05 ± 3.33 | 48 |
| Keratin 9 | KRT9 | 2.62 ± 2.83 | 17 |

Desmosome and hemidesmosome proteins increased in both porcine and human basal RPE exosomes are indicated by gray background (compare **Tables 1** and **3**). ± = SEM; \* = Abundance in exosomes released from High genetic AMD-risk iPSC lines, proteins ranked from highest to lowest.

Concomitant with the increased amounts of desmosomal and hemidesmosomal proteins in exosomes released under oxidative stress conditions, immunoblotting analyses of RPE cell lysates revealed a decrease in ITGB1 and KRT10 in the RPE lysates treated for 4 weeks with 0.2 mM H_2_O_2_ compared to controls (**Figs. 4C-D**). These complementary results further underscore that the exosomal pathway is a route of protein shedding under these chronic oxidative stress conditions.

The protein cargo identified by proteomic analysis of the apical exosomes did not reveal any obvious pathways affected by H_2_O_2_ treatments, including no changes in desmosome pathway components. This was confirmed by in depth biostatistical analyses of the apical proteomic data from the primary pig RPE cultures. As expected, no changes to desmosome shedding were found in either of these apical datasets. The only statistically significant changes found using this analysis method were a decrease in Alpha-2-Macroglobulin (A2M), Fibrillin 1 (FBN1), and Talin 1 (TLN1) in response to H_2_O_2_ treatment, (**Supp. Table S6**).

### Chronic subtoxic oxidative stress increases deposition of exosomal proteins in the ECM

We next tested whether the proteins that were increased in basolaterally released exosomes following chronic oxidative stress were also deposited in the basal ECM. We developed a protocol to lyse and remove the overlying cells, followed by rigorous solubilization of the ECM. The ECM preparations were analyzed by immunoblotting for several of the proteins increased in the basolaterally released exosomes in response to oxidative stress.

Interestingly, a subset of the basolateral desmosome and hemidesmosome exosome proteins that were increased during stress conditions (DSG1, ITGB1, and KRT10) were also significantly increased in the ECM (**Fig. 5**). Further studies will be needed to investigate whether ECM deposition of these exosome-associated proteins play a role in overall deposit formation.

**Figure 5.**
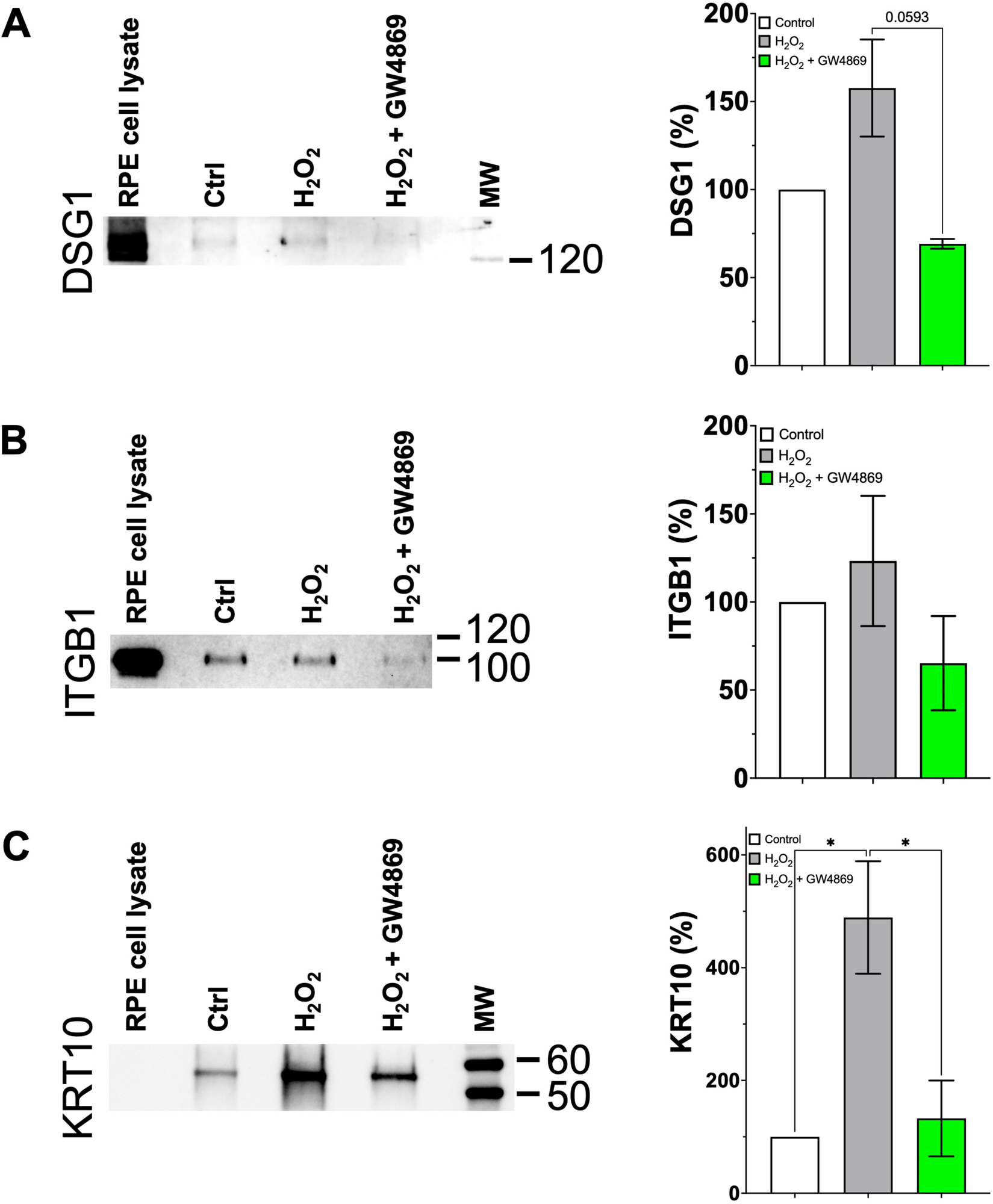
Desmosomal and hemidesmosomal proteins are deposited into the basal ECM under subtoxic oxidative stress conditions and this deposition is blocked by inhibiting exosome release. Increased deposition of DSG1 (**A**), ITGB1 (**B**), and KRT10 (**C**) into ECM is seen under oxidative stress conditions. The increased ECM deposition under oxidative stress conditions (0.2 mM H_2_O_2_) can be blocked by inhibiting exosome release with 4µM of the nSMase 2 inhibitor GW4869. Quantitation of chemiluminescent immunodetection signal is shown in the graphs on the right-hand side and are based on two to three experiments in all cases. Error bars are SEM. Data were analyzed by One-way ANOVA for treatment condition followed by Tukey’s post-hoc test for multiple comparisons. P-values below 0.1 are indicated. * = p < 0.05

### Many common drusen and ECM proteins are found in RPE exosomes, and canonical exosome proteins are found in both *ex vivo* and *in vitro* RPE ECM

We recently generated extensive proteomic data of human and porcine BrM, and porcine RPE ECM (some of the data published in (62)) and compared those data to our previously published porcine RPE basal exosomes (12). Interestingly, we found many common drusen and ECM proteins that are present in highly purified RPE exosomes. For ease of presentation and for the focus of the current study, we show a comparison of a subset of known protein components of drusen, ECM, and basally secreted exosomes in **Supp. Table S7**. The table also clearly shows the presence of a number of exosomal marker proteins in both porcine and human *ex vivo* ECM, known as Bruch’s Membrane (BrM); as well as *in vitro* RPE ECM. Finally, the table shows a high degree of similarity in the composition of human and porcine-derived samples overall, further validating the utility of the porcine model.

### Inhibition of basolateral exosome release during chronic oxidative stress decreases deposition of exosomal proteins in ECM

After establishing that some proteins found in basally secreted exosomes were also increased in ECM, we tested if inhibition of exosome release could prevent this protein accumulation in the ECM. To decrease exosome release, several different concentrations of the neutral Sphingomyelinase 2 (nSMase 2) inhibitor GW4869 were used since it has previously been shown to decrease small EV secretion (63, 64). Based on these studies we tested 4 and 20 µM concentrations and found that basolateral exosome release was significantly decreased over a 4-week treatment in fetal human RPE (**Supp. Fig. S4**). However, 20 µM caused some morphological changes in porcine RPE (**Supp. Fig. S5A**), and TER was significantly affected throughout the experiment with a near loss of barrier integrity at 4 weeks (**Supp. Fig. S5B**). Thus, we settled on 4 µM which was well tolerated morphologically (**Supp. Fig. S6A**) and caused no decrease in TER throughout the 4-week experiment (**Supp. Fig. S6B**).

Four µM GW4869 treatment concurrent with H_2_O_2_ stress prevented the increased deposition of DSG1, ITGB1, and KRT10 seen in response to H_2_O_2_ stress alone (**Figs. 5A-C**), demonstrating that this deposition is exosome-mediated.

## Discussion

In the current study, we show for the first time that chronic low-level oxidative stress in highly polarized primary RPE cultures induces basal-side specific desmosome and hemidesmosome shedding via exosomes. We also show that release of these exosomes correlates with deposition of desmosomal and hemidesmosomal proteins into ECM, and that this deposition can be prevented by inhibition of exosome release. Dysfunction of desmosomes and hemidesmosomes leads to impaired oBRB function, which is known to play a role in AMD pathology (22). The implications of oBRB breakdown in disease are profound as it results in the leakage of blood contents and influx of osmolytes that lead to accumulating fluid in the subretinal space (edema) and ultimately to exudative retinal detachment (65). Thus, early detection of RPE dysfunction is essential for efficacious intervention to maintain oBRB integrity. Significant in this regard, our findings provide potential novel biomarkers of early cellular dysfunction and opportunity for therapeutic intervention in aging-related retinal diseases, such as AMD, and broadly from blood-CNS barriers in other neurodegenerative diseases.

We investigated changes to the protein content of EVs and in particular exosomes, released both apically and basolaterally from a highly differentiated and polarized cell culture model of the outer blood-retinal barrier, under conditions of chronic subtoxic oxidative stress. Chronic subtoxic stress in the RPE (defined here as not causing overt morphological or functional dysfunction) is thought to play a central role in the pathophysiology of AMD (1) and we therefore sought to identify protein changes in RPE-derived exosomes that precede overt morphological changes and severe cellular dysregulation, in order to identify potential pre-symptomatic biomarkers of AMD.

We have previously shown that both apical and basolateral exosomes can be isolated at a high level of purity from polarized RPE cultures, and that their proteomes are distinctly different, reflecting polarized cargo loading (12). In order to adhere to the guidelines for minimum requirements for characterization of EVs that were recently described by the EV community (MISEV2018; (36)), we analyzed isolated EVs for the presence of several EV marker proteins (Syntenin, ANXA2) and other membrane-associated proteins (ITGB1), as well as the absence of markers of cellular contamination (*e.g.* Calreticulin; ER marker) by immunoblotting. We also measured the size distribution of the EV preparations, which were of expected sizes for small EVs. We implemented an improved cushioned density-gradient ultracentrifugation (C-DGUC) method for EV isolation (**Fig. 1**) that resulted in a robust enrichment of EVs while maintaining the efficient removal of contaminants seen in traditional DGUC. Analysis of the proteomes of the EV preparations clearly demonstrated that the majority of the contents were EVs, given the high percentage of membrane and transmembrane proteins, as well as canonical small EV/exosome proteins, **Supp. Tables S8-10** (basolateral EVs weeks 3&4) and **S11-12** (apical EVs weeks 3&4).

There was some variation between experiments, in which increases in exosomal basolateral-specific desmosome and hemidesmosome shedding was not only seen during weeks 3 & 4 but could also be seen in the exosomes released during weeks 1&2 (**Figs. 4A-B**). This is not surprising given that the level of resistance to stress in a subtoxic chronic setting may vary between batches of primary RPE cultures. It will be interesting to investigate in future studies in iPSC-derived RPE cultures, if this variation in stress resistance is also seen between individual donor cell lines, and if it is seen with more targeted oxidative stressors such as the mitochondrion redox cycler Paraquat.

The identification of exosome-mediated desmosomal and hemidesmosomal shedding under chronic subtoxic oxidative stress conditions, is a novel finding. Effects of oxidative stress in the RPE accumulate during aging, resulting in disruption of the outer retinal blood-barrier (66). Tight junctions are critical in maintaining this barrier and the integrity of the RPE, but how tight junctions or other structural junctions are dismantled during oxidative stress is unknown. Desmosomes are different from tight and adherens junctions, located basally to those junctions in the lateral membrane, and are specialized and highly ordered membrane domains that mediate cell-cell contact and strong adhesion, **Supp. Fig. S7** and (38). Adhesive interactions at the desmosome are coupled to the intermediate filament cytoskeleton. By mediating both cell-cell adhesion and cytoskeletal linkages, desmosomes mechanically bind cells within tissues and thereby function to resist mechanical stress (38). Hemidesmosomes on the other hand, are multiprotein complexes that facilitate the stable adhesion of basal epithelial cells to the underlying basement membrane (39), see **Supp. Fig. S7**. Strikingly, we found that desmosome- and hemidesmosome-associated proteins were being released as exosome cargo on the basolateral side, but not the apical side of the RPE. This polarized specificity is perhaps not surprising given the lateral and basal locations of desmosomes and hemidesmosomes, respectively. Nonetheless, these findings further underscore the highly polarized endosomal pathways that govern cargo loading of EVs inside the fully differentiated RPE cell. The details of how this polarized sorting of apical vs basolateral cargo into exosomes and other EVs is achieved, is not currently well understood and warrants further investigation. The current study does not distinguish between desmosomes or hemidesmosomes as the major source of the anchoring junctions shed via exosomes since proteins from both were detected. Both desmosomal (DSG1, JUP, and DSP) and hemidesmosomal (ITGB1 and ITGA3) component proteins were increased in basolaterally released exosomes under stress conditions, suggesting that they were both shed. Future studies aimed at co-immunoprecipitation experiments and similar approaches will be required to specifically clarify the relative contribution and time-course of exosome-mediated desmosomal vs hemidesmosomal shedding under chronic subtoxic oxidative stress conditions.

We are confident that we have demonstrated that desmosomes are associated with exosomes, as opposed to being shed and present as separate multiprotein complexes from exosomes, based on the following lines of evidence: 1) Desmosomes were identified in density gradient fractions containing exosomes and EVs, with densities that indicate lipid content found only in EVs and lipoprotein particles (mainly LDL; LPs); (2) In order for desmosomes to be exclusively present separate from lipid-containing vesicles and/or particles, their density (proteins have higher density than lipids) would place them in fractions with higher densities than those in which the EVs are located, and those higher density fractions were not used in our analyses; (3) Desmosomes and hemidesmosomes are plasma membrane-integral and -binding structures that are unlikely to exist separate from lipid membranes; and (4) The lipid and cholesterol content of LPs are not in the form of lipid bilayers required for intact assembly of desmosomes.

It is remarkable that highly similar protein signatures reflecting desmosome and hemidesmosome shedding were also found in basal exosomes from high AMD-risk patient-derived iPSC-RPE lines <u>not</u> subjected to oxidative stress (**Table 3** and **Supp. Table S5**). This suggests that robust common pathways in oxidative stress and AMD pathogenesis, such as gene networks that respond to environmental stressors (67), are responsible for these changes in RPE monolayers in both porcine and human cell culture models. Future studies investigating the protein content of apical and basolateral exosomes from hiPSC-RPE generated by additional laboratories and patient groups, would be important to validate the findings presented here. In addition, the use of other types of oxidative stressors, such as targeted mitochondrial uncouplers, or oxidized lipids, may help clarify the exact intracellular dysfunction causing this early-stage desmosome and hemidesmosome dismantling, and how it may affect intercellular desmosome-mediated signaling.

Oxidative stressor experiments in cell culture often raise concerns of the contributions of apoptosis and other severely destructive impacts on the cell that can lead to release of large amounts of membrane blebs. Therefore, we chose to use low levels of the oxidative stressor H_2_O_2_ over an extended timeframe in the current study. Very few previous studies have investigated the impact of prolonged oxidative stress in RPE for more than 72 hours (29) and, to our knowledge, none at subtoxic levels as in the current study. In addition, none have done so with highly purified and validated exosome isolation methods (12) as in the current study. Given the lack of any observable cell loss in light microscopic analyses, and lack of LDH release under these oxidative stress conditions (**Supp. Figs. S1A-B**), there is no reason to expect that any apoptotic processes were underway, or any release of apoptotic bodies. Furthermore, our exosome/small EV isolation protocol includes a pre-clearing centrifugation step at 10,000 x *g* for 30min prior to using the cleared supernatant for the subsequent C-DGUC steps. Thus, if any traditional apoptotic bodies (ø≈0.5-2µm; (68)) were released, this pre-clearing step would pellet and remove the vast majority of them. If a subpopulation of apoptotic bodies escapes pelleting at 10,000 x *g*, they would however locate to heavier density fractions (ϱ =1.12-1.23 g/ml) in the density gradient (69) than those used for analyses in the current study. In addition, our in-depth proteomic analyses (**Supp. Tables S8-12**) did not show an increase in traditional apoptotic body markers such as Caspases, Lamins, and ER proteins such as Calreticulin (69), in our exosome preparations under stress conditions, again demonstrating that our small EV preparations contain minimal to no amounts of apoptotic bodies.

To further ensure that our oxidative stressor conditions were indeed subtoxic, we analyzed the protein levels of the canonical RPE protein RPE65. RPE65, is an isomerase involved in the visual cycle, responsible for converting all-*trans*-retinyl ester to 11-*cis*-retinol. We did not detect a difference in RPE65 expression between control RPE cells and stressed RPE cells (**Fig. 2B**) after 4 weeks of oxidative stress. A previous study showed decreased RPE65 expression in response to H_2_O_2_-induced acute oxidative stress in RPE cultures (70). We also analyzed the levels of a tight junction protein, Occludin, which is important to maintain the RPE barrier. We did not observe a change in Occludin expression between control and stressed RPE cells (**Fig. 2C**), and this agreed with our TER data. A previous study in a porcine intestinal epithelial cell line (IPEC-J2) exposed to short-term acute H_2_O_2_-mediated oxidative stress, showed significant reduction in the protein levels of the tight junction proteins Claudin-1 and Occludin, as well as the tight junction adaptor and adherens junction protein ZO-1 (71). Similarly, RPE oxidative stress studies using 30 min to 24-hour treatment experiments with a high concentration of H_2_O_2_ (0.5-2 mM), led to loss of barrier markers such as Occludin and ZO-1 (72, 73). Thus, our lack of observed change in the RPE65 and Occludin levels demonstrate that the chronic oxidative stress conditions used in the current study were subtoxic.

Interestingly, Apolipoprotein E (ApoE), which is a common constituent of drusen in AMD eyes (74), was not increased in basolaterally released exosomes under these oxidative stress conditions (**Supp. Tables S8-10**). This is not a surprise given that ApoE is a major component of lipoprotein particles, rather than EVs released from RPE, and is yet another indicator of the purity of our exosome preparations. A recent study (29) subjecting iPSC-RPE to stress with a Cigarette Smoke Extract (CSE; (75)), reported an increase of ApoE in EVs. However, EVs in that study were isolated by differential ultracentrifugation (DUC) rather than by density gradient ultracentrifugation (DGUC) in the current study, which is considered the gold standard in the field (36). It has been shown in numerous studies (76, 77) that DUC EV preparations contain markedly higher amounts of lipoproteins compared to DGUC EV preparations. Another difference between the studies is that although both employed a 4-week treatment duration, the CSE study added stressor every 48 hours, while in the current study, the stressor (0.2mM H_2_O_2_) was added every 24 hours. Thus, the differences in choice of oxidative stressor, study design, and in EV isolation method, are likely explanations for the differing results in the two studies.

In the current study, we also showed that DSG1, ITGB1 and KRT10 are deposited in increased amounts in ECM under chronic oxidative stress conditions. Importantly, this increased deposition could be blocked by inhibiting exosome release, demonstrating the exosomal origin of these proteins. Since formation of drusen and other deposits in the BrM is a major part of the AMD disease process (1), and our data show that the exosomal pathway may play a role in this process; targeting exosome release from RPE may be feasible as a novel approach for therapeutic intervention in AMD and other retinal degenerative disorders. Different subpopulations of EVs are thought to be generated via ESCRT-dependent and ceramide-dependent (nSMase 2) biogenesis pathways, and these may be differentially utilized by apical vs basolateral EV release in highly polarized cell types (13). However, this has not been studied in detail in RPE cells to date. Furthermore, it is not clear whether inhibition of basolateral EV release will shift release of EVs to the apical side or result in rerouting of EVs to lysosomal degradation. Several studies have recently reported that increased ceramide production in RPE cells is responsible for increased RPE dysfunction and is thought to play a role in the AMD disease process (78, 79). It is not currently clear if this pathological effect is mainly or in part mediated via exosomes/EVs. Great care will be needed in choosing targets for inhibition of basolateral EV release without affecting apical release, while also avoiding disturbing endosomal trafficking overall in RPE cells. Thus, additional future studies are warranted for detailed characterization of the apical vs basolateral EV biogenesis and release pathways in RPE cells.

In conclusion, the disassembly of desmosomes and hemidesmosomes in RPE and release via exosomes during chronic oxidative stress conditions has to our knowledge not been reported previously in any epithelial cell culture system. Thus, this finding presents novel targets for interventional treatments aimed at maintaining oBRB integrity in pre-symptomatic and early-stage AMD, as well as potentially in other retinal neurodegenerative diseases. Finally, the identified chronic oxidative stress exosome protein profiles unique to RPE cells (constituting the oBRB) <u>prior</u> to overt functional and morphological dysfunction, suggest they may be candidates for early-stage AMD or even pre-AMD biomarkers accessible in the systemic circulation.

## Acknowledgments

This study was supported by NIH grants R21EY033057 (M.K.), F31EY033170 (B.J.H.), R01EY031748 (C.B.R.), R01EY022359 and R01EY028608 (W.D.S.), a grant from the Pro Retina Foundation Germany (B.H.F.W.), and a grant from the Foundation Fighting Blindness (C.B.R.). Duke University Department of Ophthalmology is supported by an unrestricted grant from Research to Prevent Blindness (RPB). In addition, a small unrestricted RPB grant to B.J.H. supported the study. A Core Grant for Vision Research (P30EY005722) from NEI (to Duke University), supported much of the work, including the mass spectrometric analyses carried out by N.P.S.

## Author Contributions

M.K., B.J.H. and N.P.S. designed and performed experiments, analyzed data, and wrote the paper; D.G., U.K. and M.A.C. designed and performed experiments, and analyzed data; K.P. and V.M. performed experiments, T.J.W. isolated primary porcine RPE cells; K.P. and B.H.F.W. provided conditioned growth media from iPSC-derived RPE cultures; A.M. and S.S.M. provided fetal RPE cells; M.S. and A.A.K. analyzed data; W.D.S. and C.B.R. designed experiments, analyzed data and wrote the paper. All authors reviewed the manuscript and agreed to its finalized version.

