## Supporting information for "Polarized Desmosome and Hemidesmosome Shedding via Exosomes is an Early Indicator of Outer Blood-Retina Barrier Dysfunction"

### Supporting Information to Hernandez *et al.* 2023

**Supporting Table S1.** Modal and average EV sizes in basolateral conditioned media from control and $H_2O_2$-treated pRPE cultures.

| | Wks 1 & 2 | | Wks 3 & 4 | |
|---|---|---|---|---|
| | **Control** | **$H_2O_2$** | **Control** | **$H_2O_2$** |
| **Modal (nm)** | 142.6 ± 13.2 | 154.6 ± 26.0 | 125.3 ± 23.6 | 143.2 ± 24.8 |
| **Average (nm)** | 144.2 ± 5.0 | 149.0 ± 8.1 | 139.5 ± 1.2 | 142.7 ± 2.7 |

Errors shown are ± SD. No statistically significant interactions or differences were detected between treatment conditions or timepoints by two-way ANOVA analysis followed by Sidak post-hoc multiple comparison analysis.

**Supporting Table S2.** Modal and average EV sizes in basolateral conditioned media from control and GW4869-treated primary human RPE cultures.

| | **Control** | **4μM GW4869** | **20μM GW4869** |
|---|---|---|---|
| **Modal (nm)** | 125.0 ± 15.1 | 119.5 ± 7.4 | 113.2 ± 6.0 |
| **Average (nm)** | 129.9 ± 6.9 | 139.4 ± 1.8 | 130.6 ± 1.8 |

Errors shown are ± SD. No statistically significant differences were detected between treatment conditions by one-way ANOVA analysis followed by Tukey post-hoc multiple comparison analysis.

**Supporting Table S3.** Proteins decreased ≥2-fold in at least two of the three sets of basolateral exosomes released during weeks 3 & 4 from $H_2O_2$-treated RPE cultures compared to untreated. Sets #1 & #2 represent two biological replicates. Set #3 was generated from exosomes released during an entire 4-week experiment. Each set was generated from two technical mass spectrometric replicate runs.

| **Protein name** | **Gene name** | **Average fold decrease ($H_2O_2$/Ctrl)** | **Average abundance ranking in $H_2O_2$ EXOs** |
|---|---|---|---|
| FXYD Domain Containing Ion Transport Regulator 1 | FXYD1 | 6.26 ± 6.08 | 109 |
| Cysteine Rich Transmembrane Module Containing 1 | CYSTM1 | 4.24 ± 1.69 | 147 |
| Coagulation Factor V | F5 | 2.49 ± 0.45 | 161 |

Errors shown are ± SEM. * = Abundance in exosomes released during $H_2O_2$ treatment, proteins ranked from highest to lowest.

**Supporting Table S4.** Proteins decreased ≥2-fold in both sets of apical exosomes released during weeks 3 & 4 from $H_2O_2$-treated RPE cultures compared to untreated. Sets #1 & #2 represent two biological replicates. Each set was generated from two technical mass spectrometric replicate runs.

| Protein name | Gene name | Average fold decrease ($H_2O_2$/Ctrl) | Average abundance ranking in $H_2O_2$ EXOs* |
|---|---|---|---|
| Galectin 3 | LGALS3 | 18.69 ± 15.55 | 612 |
| Semaphorin 3B | SEMA3B | 15.66 ± 14.78 | 630 |
| Collagen Type V Alpha 1 Chain | COL5A1 | 12.51 ± 11.22 | 532 |
| Complement Factor H | CFH | 9.92 ± 9.88 | 398 |
| ABI Family Member 3 Binding Protein | ABI3BP | 7.93 ± 1.62 | 262 |
| ATPase Plasma Membrane Ca2+ Transporting 4 | ATP2B4 | 7.38 ± 2.67 | 413 |
| Serpin Family E Member 1 | SERPINE1 | 6.28 ± 5.10 | 578 |
| Solute Carrier Family 6 Member 20 | SLC6A20 | 6.14 ± 4.82 | 401 |
| Protein Tyrosine Phosphatase Receptor Type 1 | PTPRJ | 6.03 ± 4.01 | 411 |
| Alpha-2-Macroglobulin | A2M | 6.02 ± 2.10 | 29 |
| PZP Alpha-2-Macroglobulin Like | PZP | 5.97 ± 1.05 | 99 |
| Solute Carrier Family 25 Member 3 | SLC25A3 | 5.93 ± 5.59 | 430 |
| Actinin Alpha 1 | ACTN1 | 5.51 ± 2.40 | 363 |
| H2B Clustered Histone 11 | H2BC11 | 5.35 ± 0.44 | 174 |
| Annexin A8 | ANXA8 | 5.06 ± 2.70 | 488 |
| Melanotransferrin | MELTF | 4.31 ± 1.12 | 441 |
| FERM Domain Containing Kindlin 2 | FERMT2 | 4.23 ± 1.57 | 564 |
| Teneurin Transmembrane Protein 4 | TENM4 | 4.21 ± 1.83 | 557 |
| Ribosomal Protein L3 | RPL3 | 3.83 ± 3.18 | 418 |
| Triosephosphate Isomerase 1 | TPI1 | 3.81 ± 2.49 | 304 |
| EH Domain Containing 1 | EHD1 | 3.57 ± 1.03 | 421 |
| ATPase Plasma Membrane Ca2+ Transporting 2 | ATP2B2 | 3.33 ± 0.72 | 289 |
| Growth Arrest Specific 6 | GAS6 | 3.32 ± 1.97 | 525 |
| Epidermal Growth Factor | EGF | 3.31 ± 0.33 | 281 |
| Tenascin C | TNC | 3.28 ± 0.61 | 371 |
| Dishevelled Associated Activator of Morphogenesis 1 | DAAM1 | 3.20 ± 0.80 | 527 |
| Ribosomal Protein L4 | RPL4 | 3.17 ± 1.09 | 330 |
| Heparan Sulfate Proteoglycan 2 | HSPG2 | 3.15 ± 1.01 | 471 |
| Basigin | BSG | 3.14 ± 1.16 | 97 |
| ATP Syntase F1 Subunit Alpha | ATP5F1A | 3.11 ± 0.14 | 378 |
| Formin Like 2 | FMNL2 | 3.08 ± 0.34 | 436 |
| TIMP Metallopeptidase Inhibitor 3 | TIMP3 | 3.02 ± 0.94 | 311 |
| CD151 Molecule (Raph Blood Group) | CD151 | 2.95 ± 0.47 | 370 |
| Myosin Heavy Chain 9 | MYH9 | 2.90 ± 0.76 | 107 |
| Fascin Actin-Bundling Protein 1 | FSCN1 | 2.88 ± 0.07 | 283 |
| S100 Calcium Binding Protein A10 | S100A10 | 2.77 ± 0.58 | 431 |
| ADAM Metallopeptidase Domain 10 | ADAM10 | 2.69 ± 0.92 | 335 |
| AE Binding Protein 1 | AEBP1 | 2.65 ± 0.02 | 133 |
| ATP Synthase F1 Subunit Beta | ATP5F1B | 2.55 ± 0.20 | 302 |
| Asparagine Synthetase | ASNS | 2.55 ± 0.24 | 493 |
| Insulin Like Growth Factor 1 Receptor | IGF1R | 2.51 ± 0.13 | 354 |
| Fibronectin 1 | FN1 | 2.50 ± 0.11 | 266 |
| Integrin Subunit Alpha 3 | ITGA3 | 2.49 ± 0.49 | 137 |
| Filamin B | FLNB | 2.38 ± 0.45 | 327 |

Errors shown are ± SEM. * = Abundance in exosomes released during $H_2O_2$ treatment, proteins ranked from highest to lowest.

**Supporting Table S5.** Proteins in basolateral exosomes increased ≥2-fold in at least two of four human iPSC-derived RPE lines from donors with high genetic AMD risk compared to four iPSC-RPE lines from donors with low risk (1, 2). Data was generated from two technical mass spectrometric replicate runs.

| Protein name | Gene name | Average fold increase (High-risk/Low-risk) |
|---|---|---|
| Apolipoprotein E | APOE | 4490.20 |
| Carboxypeptidase Z | CPZ | 2401.16 |
| Zinc finger protein 304 | ZNF304 | 1142.72 |
| Apolipoprotein(A) | LPA | 887.29 |
| Cadherin EGF LAG seven-pass G-type receptor 1 | CELSR1 | 558.63 |
| Apolipoprotein D | APOD | 522.80 |
| Immunoglobulin heavy variable 5-10-1 | IGHV5-10-1 | 327.95 |
| Collagen Type XI Alpha 2 Chain | COL11A2 | 156.45 |
| Nucleosome Assembly Protein 1 Like 4 | NAP1L4 | 151.91 |
| CD151 | CD151 | 144.69 |
| Tetraspanin 11 | TSPAN11 | 141.25 |
| Family With Sequence Similarity 90 Member A1 | FAM90A1 | 135.98 |
| Dermcidin | DCD | 108.37 |
| Glyceraldehyde-3-Phosphate Dehydrogenase, Spermatogenic | GAPDHS | 92.46 |
| CASC3 Exon Junction Complex Subunit | CASC3 | 75.64 |
| Chromosome 11 Open Reading Frame 80 | C11orf80 | 73.14 |
| X-Prolyl Aminopeptidase 3 | XPNPEP3 | 73.00 |
| Joining Chain Of Multimeric IgA And IgM | JCHAIN | 64.58 |
| Keratin 34 | KRT34 | 64.47 |
| Prostaglandin F2 Receptor Inhibitor | PTGFRN | 58.94 |
| Lysine Methyltransferase 2C | KMT2C | 47.57 |
| Calmodulin-binding transcription activator 1 | CAMTA1 | 46.48 |
| Complement C1q subcomponent subunit B | C1QB | 38.31 |
| NACC Family Member 2 | NACC2 | 36.82 |
| Immunoglobulin heavy variable 3-9 | IGHV3-9 | 32.95 |
| SET Domain Bifurcated Histone Lysine Methyltransferase 1 | SETDB1 | 32.28 |
| Regulatory Factor X1 | RFX1 | 30.85 |
| Shroom Family Member 3 | SHROOM3 | 30.52 |
| Cleavage stimulation factor subunit 2 tau variant | CSTF2T | 26.59 |
| Coiled-coil domain-containing protein | CCDC42 | 24.92 |
| Complement component C9 | C9 | 21.69 |
| MORC family CW-type zinc finger protein 1 | MORC1 | 21.37 |
| Clusterin | CLU | 20.21 |
| Immunoglobulin heavy variable 3-7 | IGHV3-7 | 19.89 |
| Complement factor B | CFB | 18.40 |
| Polymeric immunoglobulin receptor | PIGR | 18.04 |
| Collagen Type XI Alpha 1 Chain | COL11A1 | 17.47 |
| C-C Motif Chemokine Ligand 21 | CCL21 | 15.39 |
| Caspase Recruitment Domain Family Member 10 | CARD10 | 15.25 |
| Junction Plakoglobin | JUP | 14.93 |
| Secreted Frizzled Related Protein 1 | SFRP1 | 13.97 |
| T-Box Transcription Factor 6 | TBX6 | 12.74 |
| HEAT Repeat Containing 3 | HEATR3 | 10.59 |
| Cathepsin L1 | CTSL | 10.47 |
| Basonuclin 2 | BNC2 | 10.44 |
| Calsyntenin 2 | CLSTN2 | 10.43 |
| Elongation factor 1-alpha 1 | EEF1A1 | 9.69 |
| Fibrinogen beta chain | FGB | 9.66 |
| Lysosome-associated membrane glycoprotein 1 | LAMP1 | 9.50 |
| Elastin | ELN | 9.23 |
| Cytidine And DCMP Deaminase Domain Containing 1 | CDADC1 | 9.08 |
| Fibronectin | FN1 | 9.07 |
| Apolipoprotein A-IV | APOA4 | 9.00 |
| Immunoglobulin kappa variable 1-17 | IGKV1-17 | 8.58 |
| Ribosomal Protein S27a | RPS27A | 8.46 |
| Zinc finger protein 70 | ZNF70 | 8.08 |
| von Willebrand factor | VWF | 8.07 |
| Zinc finger protein 835 | ZNF835 | 7.71 |
| Sushi domain-containing protein 1 | SUSD1 | 7.55 |
| Immunoglobulin kappa variable 6-21 | IGKV6-21 | 6.92 |
| Radixin | RDX | 6.69 |
| Ectonucleotide Pyrophosphatase/Phosphodiesterase 2 | ENPP2 | 6.55 |
| HEAT Repeat Containing 1 | HEATR1 | 6.52 |
| Collagen alpha-1(VIII) chain | COL8A1 | 6.51 |
| Chromosome 1 Open Reading Frame 68 | C1orf68 | 6.39 |
| Amnion Associated Transmembrane Protein | AMN | 6.37 |
| Keratin 5 | KRT5 | 6.27 |

| | | |
|---|---|---|
| Nucleolar Complex Associated 4 Homolog | NOC4L | 6.11 |
| Immunoglobulin lambda constant 2 | IGLC2 | 5.90 |
| Solute Carrier Family 7 Member 5 | SLC7A5 | 5.86 |
| Serpin Family D Member 1 | SERPIND1 | 5.85 |
| Frizzled Related Protein | FRZB | 5.80 |
| Zinc finger homeobox 4 | ZFHX4 | 5.72 |
| Immunoglobulin lambda variable 7-43 | IGLV7-43 | 5.61 |
| Fibrillin 2 | FBN2 | 5.60 |
| Complement C5 | C5 | 5.58 |
| H2A Clustered Histone 1 | H2AC1 | 5.53 |
| Plasminogen | PLG | 5.40 |
| Gelsolin | GSN | 5.31 |
| Hornerin | HRNR | 5.12 |
| Keratinocyte differentiation factor 1 | KDF1 | 5.06 |
| Inhibitor Of Growth Family Member 2 | ING2 | 5.04 |
| Cell Division Cycle 123 | CDC123 | 4.89 |
| Keratin 1 | KRT1 | 4.87 |
| Keratin 16 | KRT16 | 4.84 |
| Immunoglobulin kappa variable 1-5 | IGKV1-5 | 4.82 |
| Inter-Alpha-Trypsin Inhibitor Heavy Chain 2 | ITIH2 | 4.81 |
| Complement C1q A Chain | C1QA | 4.54 |
| Keratinocyte Proline Rich Protein | KPRP | 4.51 |
| Inter-Alpha-Trypsin Inhibitor Heavy Chain 4 | ITIH4 | 4.47 |
| Apolipoprotein C-III | APOC3 | 4.36 |
| A-Kinase Anchoring Protein 13 | AKAP13 | 4.10 |
| Mitochondrial Ribosomal Protein L151 | MRPL15 | 4.07 |
| Collagen Type XVIII Alpha 1 Chain | COL18A1 | 3.98 |
| Epstein-Barr Virus Induced 3 | EBI3 | 3.93 |
| Fibrillin 1 | FBN1 | 3.90 |
| Insulin | INS | 3.90 |
| Vitronectin | VTN | 3.79 |
| PHD Finger Protein 2 | PHF2 | 3.76 |
| Keratin 2 | KRT2 | 3.75 |
| ALG5 Dolichyl-Phosphate Beta-Glucosyltransferase | ALG5 | 3.64 |
| DEAD-Box Helicase 27 | DDX27 | 3.55 |
| Von Willebrand Factor A Domain Containing 5B1 | VWA5B1 | 3.55 |
| Protein Phosphatase 1 Regulatory Subunit 26 | PPP1R26 | 3.48 |
| Desmoplakin | DSP | 3.47 |
| NOP9 Nucleolar Protein | NOP9 | 3.45 |
| Basigin | BSG | 3.43 |
| Agrin | AGRN | 3.31 |
| Inter-Alpha-Trypsin Inhibitor Heavy Chain 1 | ITIH1 | 3.23 |
| ATPase Na+/K+ Transporting Subunit Alpha 1 | ATP1A1 | 3.23 |
| CD5 antigen-like | CD5L | 3.17 |
| Immunoglobulin Lambda Like Polypeptide 5 | IGLL5 | 3.13 |
| Fibrinogen alpha chain | FGA | 3.09 |
| Semaphorin 5A | SEMA5A | 3.08 |
| Keratin 81 | KRT81 | 3.05 |
| Insulin Like Growth Factor Binding Protein 5 | IGFBP5 | 3.04 |
| Growth Arrest Specific 2 Like 1 | GAS2L1 | 3.04 |
| Solute Carrier Family 2 Member 1 (GLUT-1) | SLC2A1 | 3.00 |
| Hemoglobin subunit beta | HBB | 2.98 |
| Cadherin 17 | CDH17 | 2.94 |
| Afamin | AFM | 2.79 |
| Par-3 Family Cell Polarity Regulator Beta | PARD3B | 2.79 |
| Serpin Family A Member 1 | SERPINA1 | 2.76 |
| Cathepsin D | CTSD | 2.73 |
| Fibrinogen gamma chain | FGG | 2.72 |
| Heat Shock Protein 90 Beta Family Member 1 | HSP90B1 | 2.68 |
| Fibronectin Type III Domain Containing 1 | FNDC1 | 2.65 |
| Immunoglobulin lambda-like polypeptide 1 | IGLL1 | 2.63 |
| Secreted frizzled-related protein 5 | SFRP5 | 2.63 |
| Perilipin 4 | PLIN4 | 2.63 |
| Keratin 9 | KRT9 | 2.62 |
| SPT6 Homolog, Histone Chaperone And Transcription Elongation Factor | SUPT6H | 2.59 |
| Hemoglobin subunit alpha 1 | HBA1 | 2.59 |
| Alpha-2-macroglobulin | A2M | 2.57 |
| Annexin A2 | ANXA2 | 2.56 |
| Immunoglobulin lambda variable 3-21 | IGLV3-21 | 2.51 |
| Immunoglobulin lambda variable 1-44 | IGLV1-44 | 2.49 |
| Lipase G, Endothelial Type | LIPG | 2.46 |
| Transferrin | TF | 2.46 |
| Kallistatin | SERPINA4 | 2.45 |
| Complement C3 | C3 | 2.44 |

| | | |
|---|---|---|
| Pigment epithelium-derived factor | SERPINF1 | 2.43 |
| ATP Synthase F1 Subunit Beta | ATP5F1B | 2.38 |
| Adenomatous polyposis coli protein | APC | 2.30 |
| Immunoglobulin heavy constant gamma 4 | IGHG4 | 2.29 |
| Nucleoporin 98 And 96 Precursor | NUP98 | 2.27 |
| Immunoglobulin Kappa Variable 2D-24 (Non-Functional) | IGKV2D-24 | 2.24 |
| H2B Clustered Histone 1 | H2BC1 | 2.23 |
| Apolipoprotein B | APOB | 2.21 |
| Family With Sequence Similarity 157 Member A | FAM157A | 2.20 |
| Cytochrome C Oxidase Subunit 8C | COX8C | 2.19 |
| Tudor domain-containing protein 15 | TDRD15 | 2.19 |
| Stromal Antigen 1 | STAG1 | 2.17 |
| Rap Guanine Nucleotide Exchange Factor 1 | RAPGEF1 | 2.14 |
| Myosin Heavy Chain 6 | MYH6 | 2.13 |
| TRNA Selenocysteine 1 Associated Protein 1 | TRNAU1AP | 2.12 |
| Lactate dehydrogenase B | LDHB | 2.11 |
| Cyclin B1 | CCNB1 | 2.09 |
| CD109 | CD109 | 2.09 |
| Immunoglobulin kappa variable 3D-11 | IGKV3D-11 | 2.09 |
| Immunoglobulin heavy variable 3-15 | IGHV3-15 | 2.08 |
| Complement factor I | CFI | 2.07 |
| Immunoglobulin heavy constant gamma 2 | IGHG2 | 2.06 |
| Immunoglobulin kappa variable 2-28 | IGKV2-28 | 2.04 |
| Immunoglobulin kappa constant | IGKC | 2.02 |
| Coiled-Coil And C2 Domain Containing 2B | CC2D2B | 2.01 |
| Heparan Sulfate Proteoglycan 2 | HSPG2 | 2.00 |

Proteins involved in desmosome and hemi-desmosome structure and function are indicated by gray background.

**Supporting Table S6.** Likelihood-Ratio Test was used as a quantitative measure to analyze proteins in apical exosomes released from $H_2O_2$-treated pig RPE cultures compared to untreated. The table shows the proteins that were significantly decreased in response to $H_2O_2$ treatment in two biological replicate datasets. Each set was generated from two technical mass spectrometric replicate runs. Raw abundance peptide spectral counts were normalized using DESeq2 R package (3). Combined p-values were generated from a meta-analysis of data across mass spectrometry runs with metafor (4).

| Protein name | Gene name | Average fold decrease ($H_2O_2$/Ctrl) | p-value |
|---|---|---|---|
| Alpha-2-Macroglobulin | A2M | 7.62 | $8.14 \times 10^{-14}$ |
| Fibrillin 1 | FBN1 | 4.82 | $3.85 \times 10^{-10}$ |
| Talin 1 | TLN1 | 3.23 | $9.50 \times 10^{-4}$ |

**Supporting Table S7. Common drusen and ECM proteins are found in highly purified RPE exosomes.** Proteins involved in drusen formation, ECM turnover, and the exosome pathway that were found in normal Human & Porcine BrM, Porcine RPE (pRPE) ECM, and basolateral exosomes by our recent LC-ESI-MS/MS OrbiTrap proteomic analyses.

| | | BrM | | Porcine | | |
|---|---|---|---|---|---|---|
| **Protein** | **Gene** | **Human** | **Porcine** | **ECM** | **Basal EXOs** | **Location/ Function** |
| | | UP | UP | UP | UP | |
| ***Drusen components*** | | | | | | |
| Enolase 1 | ENO1 | 30 | 6 | 7 | 6 | DRU, EXO |
| Complement C3 | C3 | 28 | 29 | 5 | 5 | DRU, AMD |
| Annexin A5 | ANXA5 | 27 | 24 | ND | 21 | DRU, EXO |
| Complement Factor H | CFH | 23 | 3 | ND | 1 | DRU, AMD |
| Annexin A2 | ANXA2 | 21 | 31 | 16 | 28 | DRU, EXO |
| Retinol Dehydrogenase 5 | RDH5 | 18 | 2 | 6 | 2 | DRU, RPE |
| Clusterin | CLU | 17 | 3 | 13 | 12 | DRU |
| Vitronectin | VTN | 14 | 6 | ND | ND | DRU |
| Amyloid P Component, Serum | APCS | 13 | 6 | ND | ND | DRU |
| Crystallin Alpha B | CRYAB | 13 | 1 | 10 | 4 | DRU, RPE, EXO |
| Complement C9 | C9 | 10 | 1 | ND | 1 | DRU |
| TIMP Metallopeptidase Inhibitor 3 | TIMP3 | 9 | ND | 17 | 5 | DRU, ECM, MD |
| Apolipoprotein E | APOE | 9 | 13 | 10 | 33 | DRU, AMD |
| Complement C5 | C5 | 9 | 11 | ND | 1 | DRU |
| Annexin A1 | ANXA1 | 8 | 14 | 9 | 17 | DRU, EXO |
| CD63 | CD63 | 6 | ND | ND | 2 | DRU, EXO |
| Glycoprotein NMB | GPNMB | 4 | ND | 6 | 2 | DRU, RPE |
| ATP Synthase F1 Subunit Beta | ATP5F1B | 2 | 17 | 13 | 10 | DRU, EXO |
| Fibronectin 1 | FN1 | 1 | 30 | 10 | 4 | DRU, ECM |
| HtrA Serine Peptidase 1 | HTRA1 | 1 | ND | 27 | 2 | DRU, ECM, AMD |
| ***ECM components*** | | | | | | |
| Heparan Sulfate Proteoglycan 2 | HSPG2 | 60 | 15 | 9 | ND | ECM |
| Keratin 1 | KRT1 | 38 | 2 | 8 | 15 | ECM |
| Keratin 2 | KRT2 | 20 | ND | 6 | 8 | ECM |
| Keratin 10 | KRT10 | 12 | ND | 10 | 15 | ECM |
| EGF-Containing Fibulin Extracellular Matrix Protein 2 | EFEMP2 | ND | 1 | 7 | ND | ECM |
| Lysyl Oxidase Like 1 | LOXL1 | ND | ND | 25 | 12 | ECM |
| EGF-Containing Fibulin Extracellular Matrix Protein 1 | EFEMP1 | ND | ND | 24 | 17 | ECM, MD |
| Fibulin 2 | FBLN2 | ND | ND | 13 | 15 | ECM |
| ADAMTS Like 5 | ADAMTSL5 | ND | ND | 18 | 1 | ECM |
| Fibrilllin 2 | FBN2 | ND | ND | 1 | ND | ECM |
| ***Exosome markers*** | | | | | | |
| Actinin Alpha 4 | ACTN4 | 41 | 45 | 17 | 17 | EXO |
| Heat Shock Protein 90 Alpha Family Class A Member 1 | HSP90AA1 | 37 | 25 | 6 | 4 | EXO |
| Heat Shock Protein Family A (Hsp70) Member 5 | HSPA5 | 28 | 24 | 9 | 5 | EXO |
| Integrin Subunit Alpha V | ITGAV | 19 | ND | 3 | 19 | EXO, RPE |
| Syntenin 1 | SDCBP1 | 2 | ND | ND | 16 | EXO |
| CD81 | CD81 | 2 | ND | ND | 8 | EXO, RPE |

UP=The unique number of peptides (UP) that were identified indicates relative quantity. ND=Not detected, DRU=Drusen component, EXO=Known exosome markers, ECM=ECM component, AMD=AMD risk association, MD=Macular Dystrophy mutation, RPE=RPE marker. Known exosome markers are shaded in gray.

**Supporting Tables S8-S10.** Excel files containing lists of all the proteins identified in three separate basolateral pig RPE exosome preparations with $H_2O_2$ to Control ratio values and abundance values. Tables S8-S9 (Sets #1 and #2) were generated from exosomes released during weeks 3 & 4. Table S10 (Set #3) was generated from exosomes released during weeks 1-4. Proteins sorted according to enrichment into the exosome preparations from $H_2O_2$-treated RPE cultures are shown in the tab “$H_2O_2$ enrichment” and proteins sorted according to abundance in the exosome preparations from $H_2O_2$-treated RPE are shown in the tab “$H_2O_2$ abundance”.

**Supporting Tables S11-S12.** Excel files containing lists of all the proteins identified in two separate preparations (Sets #1 and #2) of apical pig RPE exosomes released during weeks 3 & 4 of experiments, with $H_2O_2$ to Control ratio values and abundance values. Proteins sorted according to enrichment into the exosome preparations from $H_2O_2$-treated RPE cultures are shown in the tab “$H_2O_2$ enrichment” and proteins sorted according to abundance in the exosome preparations from $H_2O_2$-treated RPE are shown in the tab “$H_2O_2$ abundance”.

**Supporting Tables S13-S16.** Excel files containing lists of all the proteins identified in four separate preparations (Sets #1-4) of basolateral exosomes released from human iPSC-derived RPE lines from donors with high genetic AMD risk compared to four iPSC-RPE lines from donors with low risk (1, 2). High-risk lines to Low-risk lines ratio values and abundance values are included. Proteins sorted according to enrichment into the exosome preparations from High-risk RPE cultures are shown in the tab “High-risk enrichment” and proteins sorted according to abundance in the exosome preparations from $H_2O_2$-treated RPE are shown in the tab “High-risk abundance”.

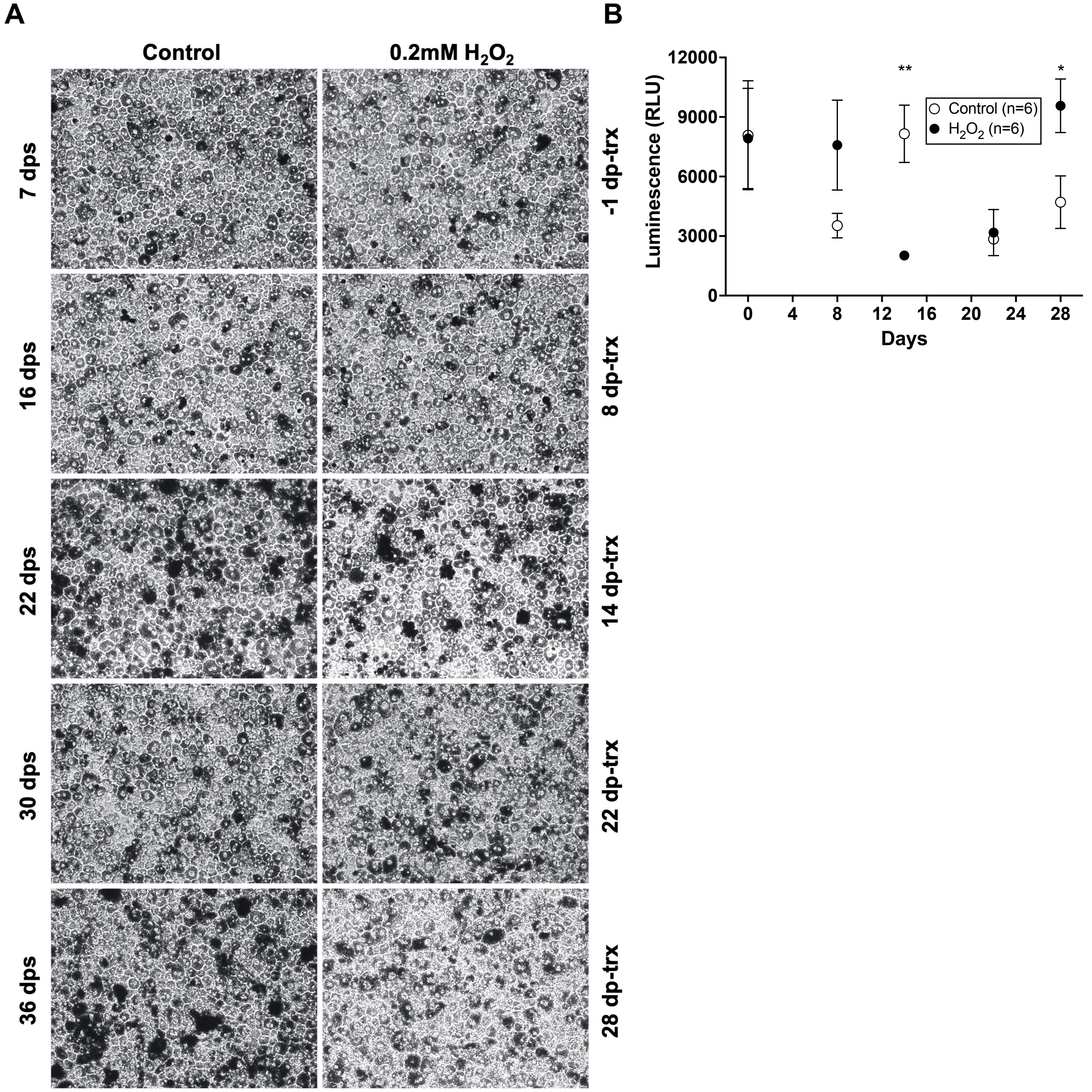


**Supporting Figure S1. Chronic subtoxic oxidative stress conditions do not affect RPE morphology or cause cytotoxicity.** (**A**) Light micrographs of pRPE transwell cultures untreated or treated with 0.2 mM $H_2O_2$. Characteristic hexagonal cell shape was maintained under this chronic $H_2O_2$ treatment condition. dps = days post seeding, dp-trx = days post start of treatment (**B**) A bioluminescent cytotoxicity assay measuring release of Lactate Dehydrogenase (LDH) in apical media. Chronic 0.2 mM $H_2O_2$ treatment did not cause significant cytotoxicity throughout the duration of treatment, further demonstrating that the chosen oxidative stress conditions were subtoxic. Error bars are SEM. Statistical analysis by multiple unpaired t-tests assuming equal variance is displayed as * = $p < 0.05$, ** = $p < 0.01$. RLU = Relative luminescence units

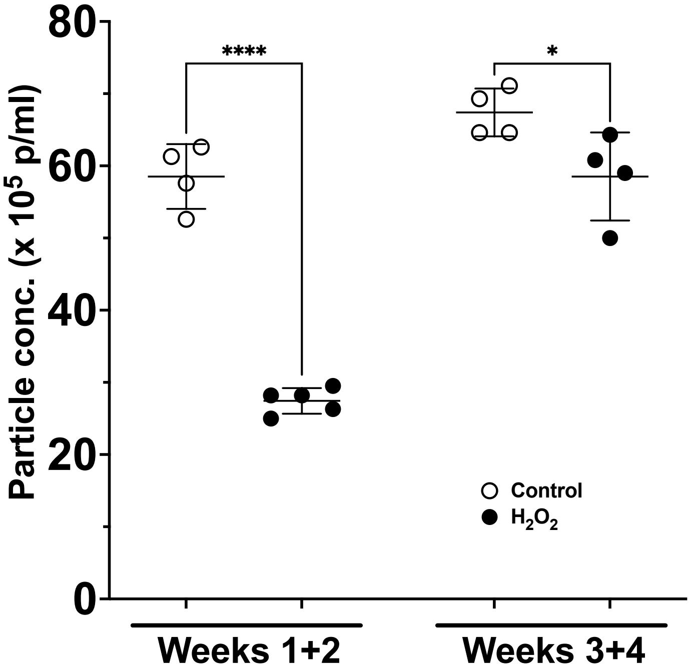


**Supporting Figure S2. Nanoparticle tracking analysis (NTA) to quantify the number of apically released exosomes.** Exosomes released from porcine RPE cultures showed a decrease in response to $H_2O_2$ treatment. Error bars are standard deviation. Data were analyzed by Two-way ANOVA followed by Tukey's post-hoc test. * = $p < 0.05$, **** = $p < 0.0001$

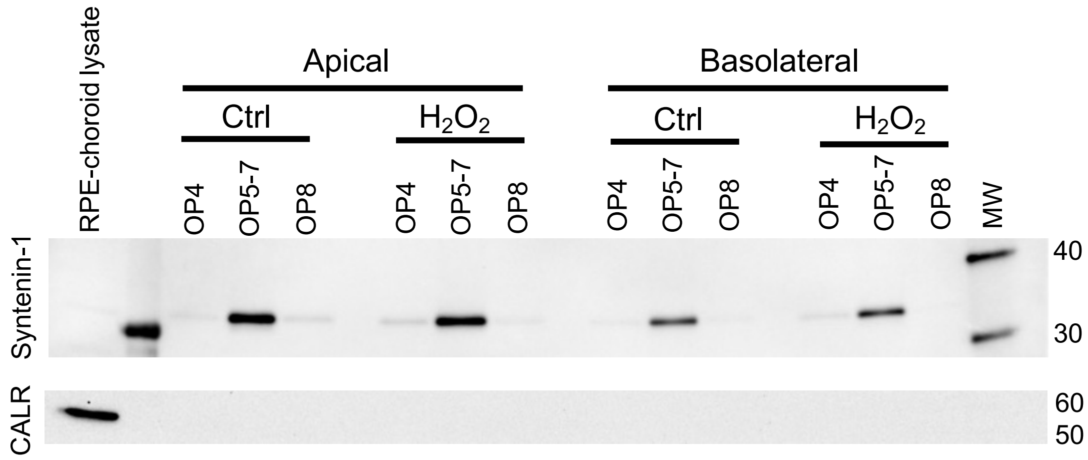


**Supporting Figure S3. Immunoblot of apical and basolateral exosomes in C-DGUC fractions from Weeks 3 & 4 of control and $H_2O_2$-treated pRPE cultures.** The canonical exosome marker Syntenin-1 is present in EVs that float at the known exosome densities of 1.07-11 g/ml in fractions 5 through 7, and only present at very low levels in lighter or heavier fractions. The ER marker Calreticulin (CALR) which is considered a marker of cellular contamination in EV preparations (5), is not present in any of these fractions. Lysate of RPE-choroid from an *ex vivo* porcine eye is included as a positive control for Syntenin-1 and CALR. Note the enrichment of Syntenin-1 in C-DGUC fractions (2µg total protein) compared to the RPE-choroid tissue lysate (10µg total protein); and conversely the reduction of Calreticulin in C-DGUC fractions compared to tissue lysate.

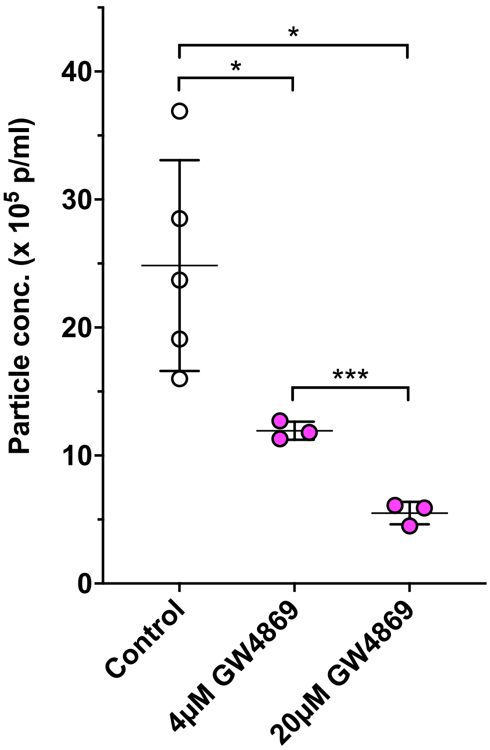


**Supporting Figure S4. NTA of basolateral exosome release in response to a Neutral Sphingomyelinase 2 (nSMase 2) inhibitor treatment.** NTA of basolaterally released exosomes from fetal human RPE cultures, showed significant decrease in response to treatment with the Neutral Sphingomyelinase 2 inhibitor GW4869. Error bars are standard deviation. Unpaired two-tailed t-test assuming equal variance was used to assess statistical significance. * = $p < 0.05$, *** = $p < 0.001$

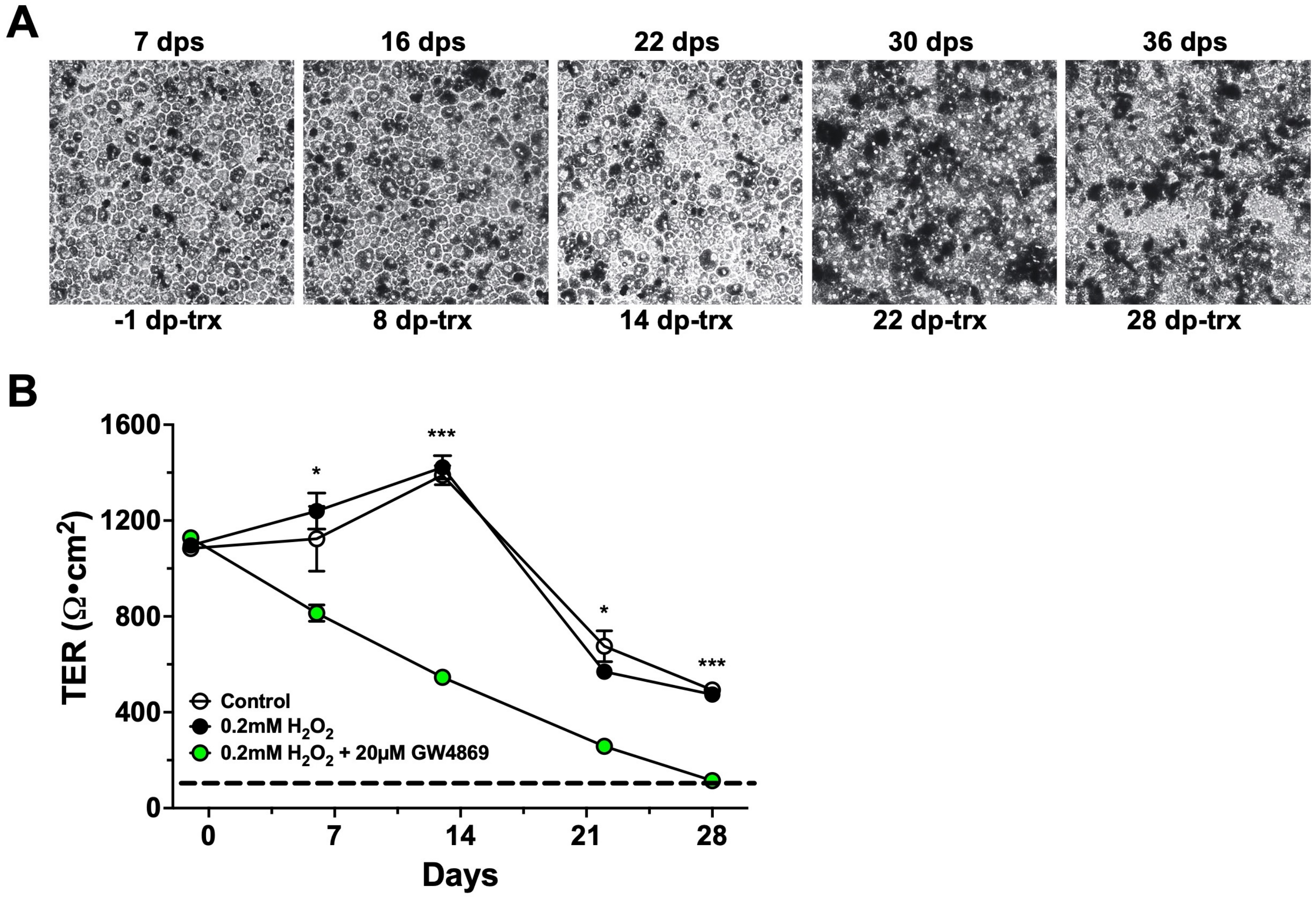


**Supporting Figure S5. Long-term treatment with a high concentration of nSMase 2 inhibitor is toxic to RPE cell cultures.** Light micrographs and TER measurements of pRPE transwell cultures treated with 0.2 mM $H_2O_2$ and 20µM GW4869. (**A**) Concurrent treatment with 0.2 mM $H_2O_2$ and 20µM of an inhibitor of nSMase2 (GW4869) increased pigmentation and caused some loss of hexagonal cell shape. dps = days post seeding, dp-trx = days post start of treatment (**B**) TER was significantly affected throughout the experiment with a near loss of barrier integrity at 4 wks for cultures treated with 0.2 mM $H_2O_2$ + 20µM GW4869. The approximate TER for a minimally intact epithelial barrier (~100 $\Omega \bullet cm^2$) is indicated with a dashed line, see (6). Error bars are SEM. Data were analyzed by Two-way ANOVA followed by Tukey's post-hoc test. The smallest statistically significant difference between $H_2O_2$+GW4869 and either of $H_2O_2$ or Control treatments, are indicated. * = $p < 0.05$, *** = $p < 0.001$

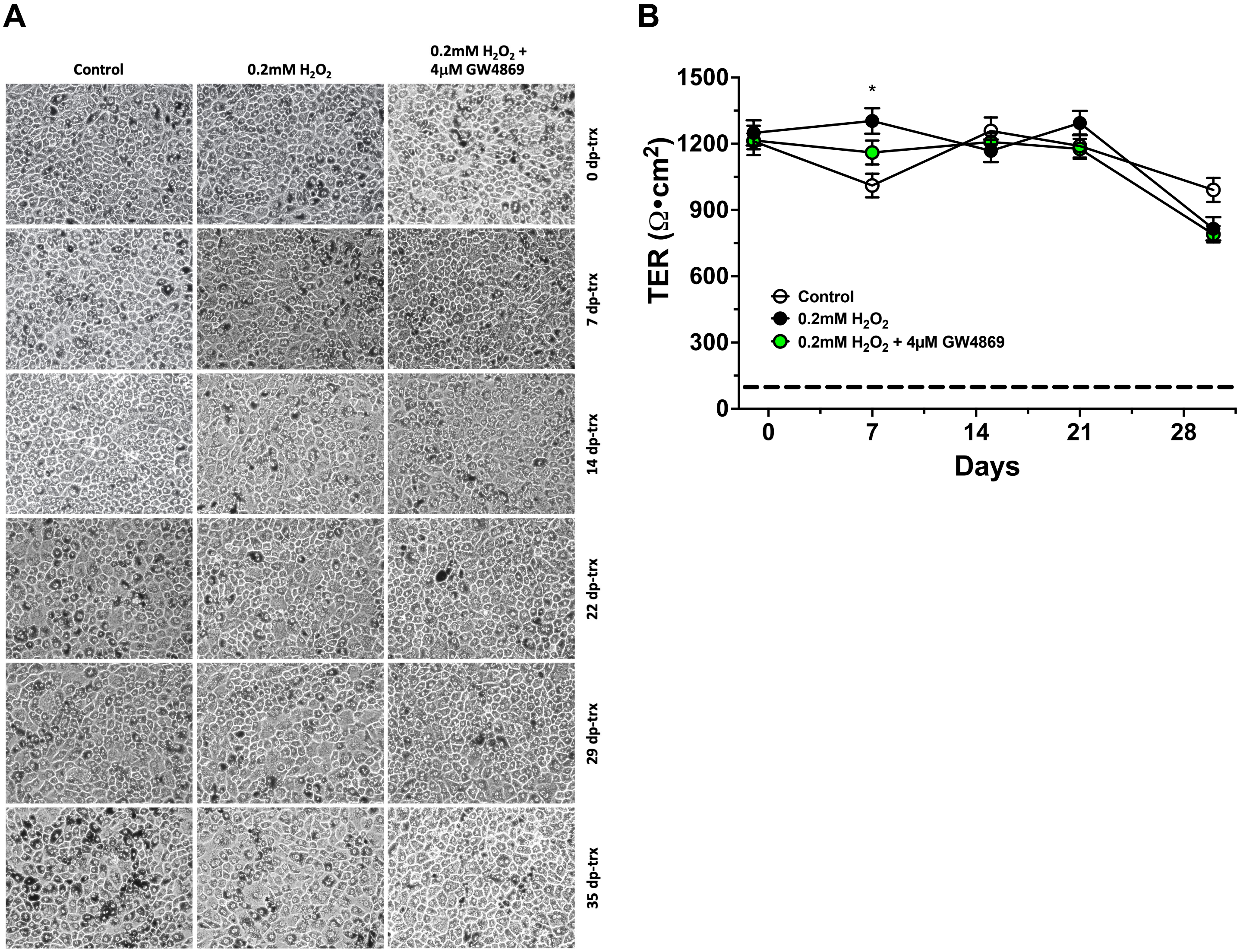


**Supporting Figure S6. Long-term treatment with 4 µM of the nSMase 2 inhibitor GW4869 is well tolerated in RPE cell cultures.** (**A**) Light micrographs of pRPE cultures in flasks untreated, treated with 0.2 mM $H_2O_2$, and treated with 0.2 mM $H_2O_2$ + 4µM GW4869. Concurrent treatment with 0.2 mM $H_2O_2$ and 4µM GW4869 did not affect apparent pigmentation levels or cause loss of hexagonal cell shape. (**B**) TER was not significantly affected in RPE cultures throughout a 4-week treatment with 0.2 mM $H_2O_2$ + 4µM GW4869. The approximate TER for a minimally intact epithelial barrier (~100 $\Omega \bullet cm^2$) is indicated with a dashed line, see (6). Error bars are SEM. Data were analyzed by Two-way ANOVA followed by Tukey's post-hoc test. Statistically significant difference between Control and $H_2O_2$ treatment is indicated as * = $p < 0.05$

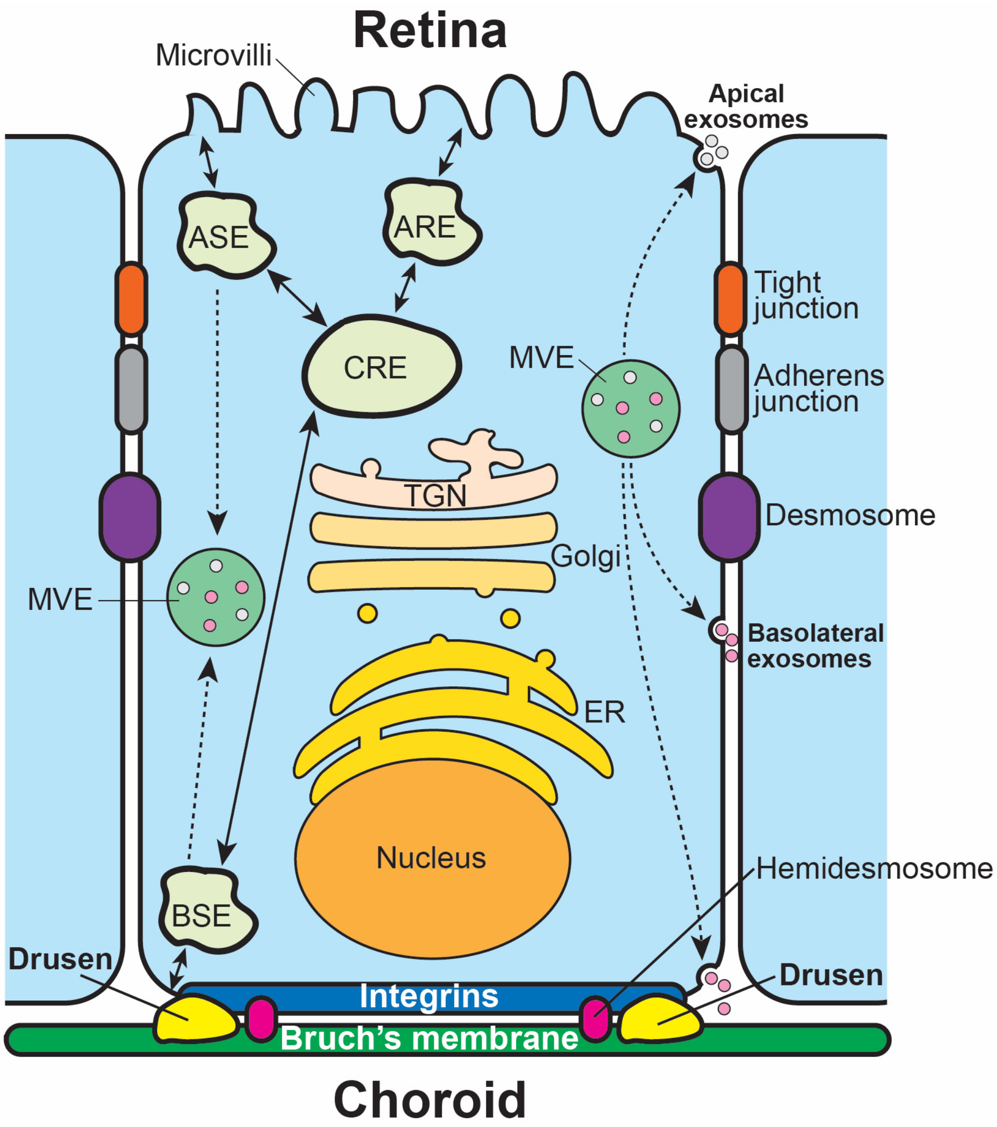


**Supporting Figure S7. Schematic of a fully differentiated retinal pigmented epithelium (RPE) cell highlighting endosomal compartments important for polarized trafficking.** The RPE cell has ciliary structures called 'microvilli' on the apical membrane surface facing the retina, which provide close physical interaction with photoreceptor outer segments (POS) and support the canonical POS phagocytosis required for renewal and recycling of POS disc components and thus functional vision. Essential for the RPE's function to provide nutrients to photoreceptors, is the transport from the choroidal vasculature (Choroid) on its basal side across the pentalaminar collagen- and elastin-rich extracellular matrix (ECM) known as Bruch's membrane (BrM), and the reverse transport of waste products back into the systemic circulation. The RPE cell monolayer in the eye serves as the outer blood-retina-barrier (oBRB), and to carry out this function RPE cells have 'tight junction' structures that control all ion flow in and out of the posterior eye. Tight junctions (orange ovals) between cells are connected areas of the plasma membrane that stitch cells together. Adherens junctions (gray ovals) join the actin filaments of neighboring cells together. Desmosomes (purple ovals) are even stronger connections that join the intermediate filaments (keratins) of neighboring cells. Hemidesmosomes (magenta ovals) connect intermediate filaments of the cell to the basal ECM, i.e., BrM. Non-polarized cells have several different endosomal compartments such as sorting endosomes (SE), and recycling endosomes (RE); as well as multivesicular endosomes (MVE) responsible for small EV and exosome biogenesis. Highly polarized cells, however, have specialized polarized endosomal compartments such as apical sorting endosomes (ASE), apical recycling endosomes (ARE), common recycling endosomes (CRE), and basal sorting endosomes (BSE). It is currently unclear how this polarized endosomal cargo sorting is maintained in exosomal cargo loading in MVEs. 'Drusen' are protein- and lipid-rich extracellular deposits that form between the basal lamina of the RPE and the inner collagenous layer of BrM. Accumulation of drusen is a common early sign of age-related macular degeneration (AMD).
